# Bone marrow hemogenic endothelial cells contribute multilineage hematopoietic progenitors in adult mice

**DOI:** 10.1101/2022.09.09.507292

**Authors:** Jing-Xin Feng, Mei-Ting Yang, Caiyi C. Li, Ferenc Livák, Abdalla Abdelmaksoud, Dunrui Wang, Giovanna Tosato

## Abstract

During development, hematopoietic stem/progenitor cells (HSPCs) originate from a subset of hemogenic endothelial cells (ECs) through a process of endothelial-to-hematopoietic transition (EHT). This process is temporally restricted to short developmental windows and generates HSPC with distinct capabilities for hematopoiesis. Although it is generally thought that adult hematopoiesis is sustained by HSCs derived from hemogenic endothelium during development, some observations point to EHT persistence in the late fetus/perinatally. Here we use lineage tracking and bioinformatics analysis to assess the presence of hemogenic endothelial cells in the adult mouse. Our analysis identifies a subset of bone marrow-resident adult endothelial cells, characterized by the expression of VE-Cadherin and the transcription factor RUNX1, that produce CD45^+^ hematopoietic cells. This EHT generates hematopoietic progenitors, and mature myeloid and lymphoid cells in the adult mouse. Our results reveal the identification of a distinct source of adult blood.

## INTRODUCTION

Hemogenic endothelial cells (ECs) generate hematopoietic cells during development through a process of endothelial-to-hematopoietic transition (EHT) at geographically defined anatomical sites (Bertrand et al., 2010; Boisset et al., 2010; Chen et al., 2011; Frame et al., 2016; Jaffredo et al., 1998; Nakano et al., 2013; Rhodes et al., 2008; Yokomizo and Dzierzak, 2010; Zovein et al., 2008). At all these locations, the hemogenic ECs represent a small fraction of all ECs (Goldie et al., 2008; Hirschi, 2012; Marcelo et al., 2013), and their competency to produce hematopoietic stem and progenitor cells is temporally restricted to short developmental windows, and their hemogenic potential differs (de Bruijn et al., 2002; Hadland et al., 2015; Iturri et al., 2021; Lin et al., 2014; Soares-da-Silva et al., 2021; Yoshimoto et al., 2012). The hemogenic endothelium of the dorsal aorta produces hematopoietic stem cells (HSC) and other multi-potent progenitors between E10.5 and E11.5 (de Bruijn *et al*., 2002; Hadland *et al*., 2015; Patel et al., 2022) that persist in the adult and are believed to sustain adult hematopoiesis throughout life. Although hemogenic ECs were thought to be restricted to the early stages of mouse development, recent experiments indicate that a hemogenic endothelium capable of multilineage hematopoiesis is also present in the bone marrow (BM) at birth and perinatally (Yvernogeau et al., 2019).

The transcription factor Runx1, which marks the hemogenic endothelium, is required for hematopoiesis from hemogenic endothelium and can confer a hemogenic potential to embryonic ECs that do not have such potential (Chen et al., 2009; Eliades et al., 2016; North et al., 1999; Yzaguirre et al., 2018). When RUNX1 was expressed in adult ECs in conjunction with the transcription factors FOSB, GFI1, and SPI1, transplantable HSC capable of BM hematopoietic reconstitution was generated (Lis et al., 2017). Nonetheless, the existence of mature ECs with persistent hemogenic potential in the adult mouse has not been demonstrated or excluded (Hirschi, 2012; Kilani et al., 2019; Pelosi et al., 2012; Zovein *et al*., 2008). Aided by technological advances in cell lineage tracking and single-cell technologies (Hernandez et al., 2022), we report the identification of a population of hemogenic ECs in the adult mouse BM that is a competent producer of hematopoietic progenitors and mature blood cells.

## RESULTS

### Adult endothelial cells generate hematopoietic cells

Since Cdh5 (codes for Vascular Endothelial Cadherin, VE-Cadherin) is selectively expressed by all ECs but not by hematopoietic cells, activation of the Cdh5-Cre^ERT2^ recombinase activity in adult mice carrying a Cre reporter is expected to track hematopoietic output from adult hemogenic ECs (Gentek et al., 2018). Therefore, we generated three Cdh5-based lineage-tracking models and activated Cre recombination in adult mice to test whether EHT is present in the adult mouse. The lineage-tracking lines were generated from the inducible Cdh5-Cre mouse lines Cdh5-Cre^ERT2^(PAC) (Pitulescu et al., 2010; Sorensen et al., 2009) and Cdh5-Cre^ERT2^(BAC) (Okabe et al., 2014), and the Cre reporter mouse lines mTmG and ZsGreen for expression of the fluorescent proteins EGFP on the cell membrane (mG) (Muzumdar et al., 2007) and ZsGreen in cell bodies (Madisen et al., 2010), respectively (Figure 1A). Four to six-week-old mice were treated with three doses of tamoxifen on consecutive days. Peripheral blood and BM were harvested four weeks after tamoxifen administration (Figure 1B). By flow cytometry, in all three mouse lines, most BM VE-Cadherin^+^CD31^+^ ECs were tracked by tamoxifen-induced fluorescence (Figure S1A and S1B). Also, by imaging, most BM Endomucin^+^ ECs showed ZsGreen intracellular fluorescence (Figure S1C and S1D). Confirming previous observations that reporter genes differ in their susceptibility to basal Cre^ERT2^ activity (Álvarez-Aznar et al., 2020), we detected a low-level tamoxifen-independent reporter fluorescence in BM ECs from the ZsGreen reporter lines (Figure S1B-S1D), which was minimal in the mTmG reporter line (Figure S1A).

**Figure 1.**
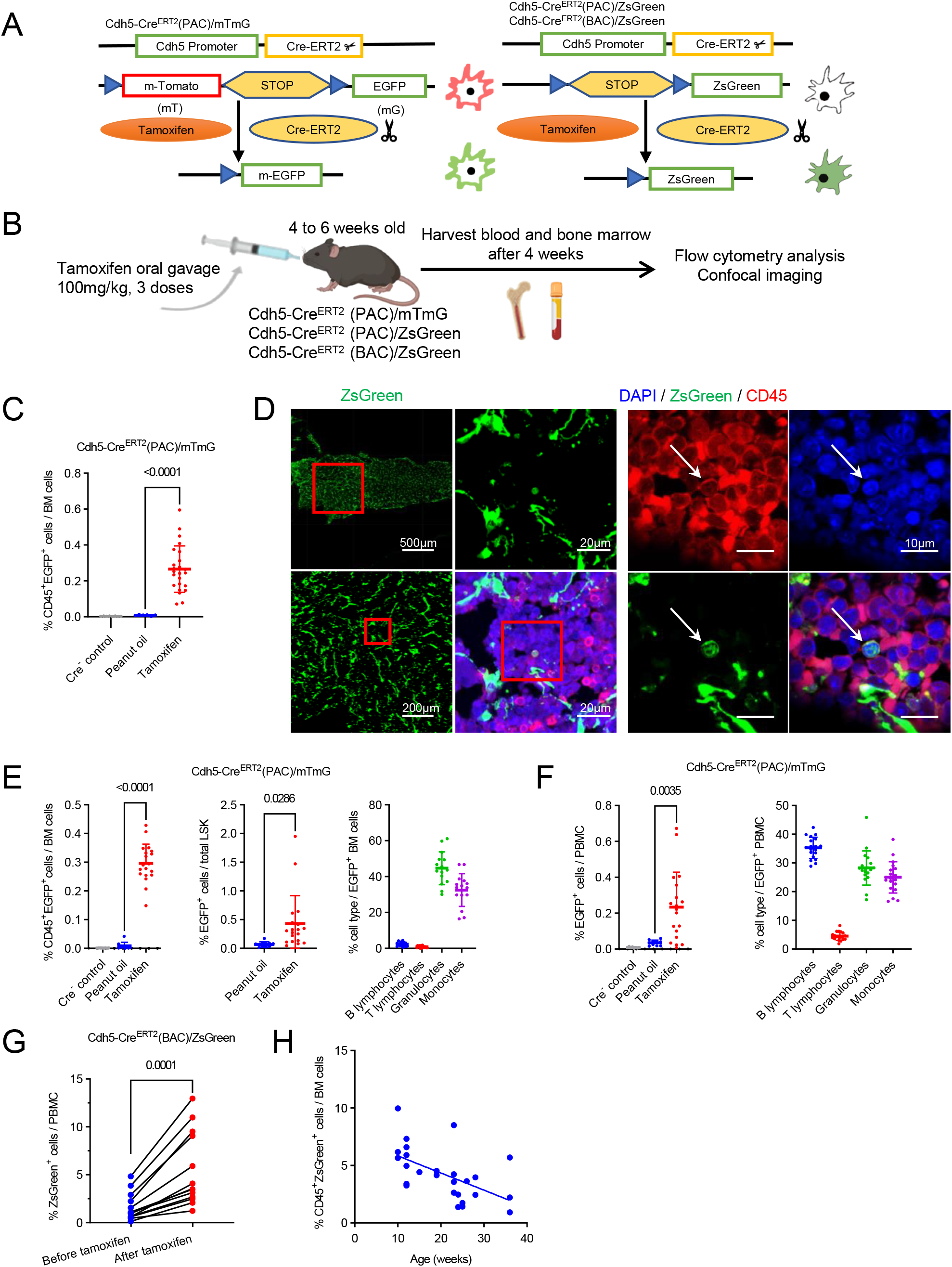
Lineage tracking discloses a contribution of endothelial cells to hematopoiesis in adult bone marrow. (A) Schematic of the Cdh5-tracking mouse lines. Tamoxifen switches-on green fluorescence in all cells that express the Cre-recombinase and their cell progeny. (B) Schematic of experiments showing the timing of tamoxifen-induced switch-on of fluorescence in VE-Cadherin (*Cdh5*)-expressing cells, tissue sampling, and analysis. (C) Percent EGFP^+^CD45^+^ cells in BM cells from Cre^-^ control, Cre^+^ control (peanut-oil treated) and Cre^+^ tamoxifen-treated mice. The horizontal line reflects the mean; each dot represents results from individual mice (1 femur plus 1 tibia combined). Statistical significance of group differences (where noted in this figure) is indicated by the *P* value (unpaired two-tailed Student’s *t*-test). (D) Representative confocal images of a BM ZsGreen^+^CD45^+^ cell from a Cdh5-Cre^ERT2^(PAC)/ZsGreen mouse. Lower magnification images show derivation of the magnified images. (E, F) Percent CD45^+^EGFP^+^ cells in BM cells (left); percent EGFP^+^ cells in BM LSK (Lin^-^ Sca1^+^cKit^+^) cells (middle); and percent B and T-lymphocytes, granulocytes, and monocytes in EGFP^+^ BM cells (right) (E). Percent EGFP^+^ cells in PBMC (left); percent B and T lymphocytes, granulocytes, and monocytes in EGFP^+^ PBMC (right) (F). Each dot represents results from individual mice (1 femur plus 1 tibia combined, or 500μL peripheral blood). (G) Percent ZsGreen^+^ cells in PBMC of individual Cdh5-Cre^ERT2^(BAC)/ZsGreen mice before or four weeks after tamoxifen administration. Each dot represents the results from 50-250μl blood/mouse. (H) Time-dependent decline of ZsGreen^+^CD45^+^ cells in BM samples from tamoxifen-treated Cdh5-Cre^ERT2^(BAC)/ZsGreen mice. Mouse age reflects age at the time of harvest. See also Figure S1 and S2.

Using these mouse lines, we examined the hemogenic potential of the tracked ECs. Flow cytometry showed the existence of fluorescence tracked CD45^+^ hematopoietic cells in the BM indicative of their endothelial derivation. Such cells were tracked in each of the three mouse lines and were significantly expanded by tamoxifen administration to adult mice (4-6 weeks old) (Figure 1C and S1E-S1G). Additionally, confocal microscopy revealed the presence of isolated, rounded CD45^+^ cells of endothelial lineage (Cdh5-tracked, ZsGreen^+^) in the BM from tamoxifen-induced adult mice (Figure 1D), suggesting that hemogenic ECs may reside in the adult BM. Characterization of these tracked BM CD45^+^ cells showed the presence of hematopoietic LSK (Lin^-^cKit^+^Sca1^+^) progenitors, CD11b^+^Ly6G^+^ granulocytes, and CD11b^+^Ly6G^-^ monocytes and fewer CD19^+^ B and CD3^+^ T lymphocytes (Figure 1E and Figure S2A). Control experiments demonstrated that tamoxifen does not alter the overall (tracked plus non-tracked cells) distribution of these hematopoietic cell subsets four weeks after administration (Figure S2B and S2C). The peripheral blood mononuclear cells (PBMC) of tamoxifen treated Cdh5-Cre^ERT2^(PAC)/mTmG and Cdh5-Cre^ERT2^(PAC)/ZsGreen adult mice also contained EGFP^+^ (Figure 1F) and ZsGreen^+^ (Figure S2D and S2E) cells, which included mature, granulocytes, monocytes, B lymphocytes and rare T lymphocytes (Figure 1F and S2E). Control experiments showed that tamoxifen does not alter the overall distribution of blood cell populations (Figure S2F and S2G).

In the ZsGreen reporter lines, we detected a fraction of fluorescent CD45^+^ cells in tamoxifen untreated Cre^+^ mice, but not in Cre^-^ mice, raising the possibility that these reporter positive CD45^+^ hematopoietic cells had been labelled earlier in development prior to tamoxifen administration, at a time when the Cdh5^+^ endothelial precursors were hemogenic (Figure S1E). However, the significant increase of tracked CD45^+^ hematopoietic cells indicates that EHT has occurred in the adult BM (Figure S1B and S1E). This is further supported by experiments showing that individual mice displayed a significant increase in the percentage of CD45^+^ZsGreen^+^ cells after treatment with tamoxifen compared to before tamoxifen treatment (Figure 1G).

Next, we examined the effect of mouse age on the endothelial hemogenic potential by treating with tamoxifen at different time points between week 6 and 30 of age and testing after 4 weeks (week 10-36). The results support the presence of CD45^+^ hematopoietic cells of endothelial derivation in the adult (24-week-old) mouse and indicate that the hemogenic potential of ECs progressively declines with age (Figure 1H).

### Single cell transcriptomic identifies a population of putative hemogenic endothelial cells in adult bone marrow

Since EHT is strictly dependent upon the transcription factor Runx1 during mouse development (Boisset *et al*., 2010; Chen *et al*., 2009; Kissa and Herbomel, 2010; Lancrin et al., 2009; North *et al*., 1999; Yokomizo et al., 2001), we sought to identify Runx1-expressing ECs in the adult mouse BM and probe their hemogenic potential. To identify this population, we mined publicly available single-cell RNAseq datasets of hematopoietic-cell depleted BM cell populations from adult mice 1 to 16 months-old (Baccin et al., 2020; Baryawno et al., 2019; Sivaraj et al., 2021; Tikhonova et al., 2019; Zhong et al., 2020) and from the caudal arteries of mouse embryos 9.5 to 11.5 days post coitum (Zhu et al., 2020), overall comprising ∼375,000 cells. Louvain clustering (Blondel et al., 2008) revealed 37 clusters (Figure S3A and S3B), and cell annotation revealed the presence of ECs, fibroblasts, adipocytes, neuronal cells, hematopoietic and other cells (Figure S3C). Clusters 15 (98% of cells derived from the embryonic dataset (Zhu *et al*., 2020)) and 26 (95.5% of cells derived from the adult datasets) (Figure S3D) were distinctive from all other clusters in harboring cells co-expressing Cdh5 and Runx1 (Figure 2A and 2B), and showing a close relationship in gene (Figure 2C).

**Figure 2.**
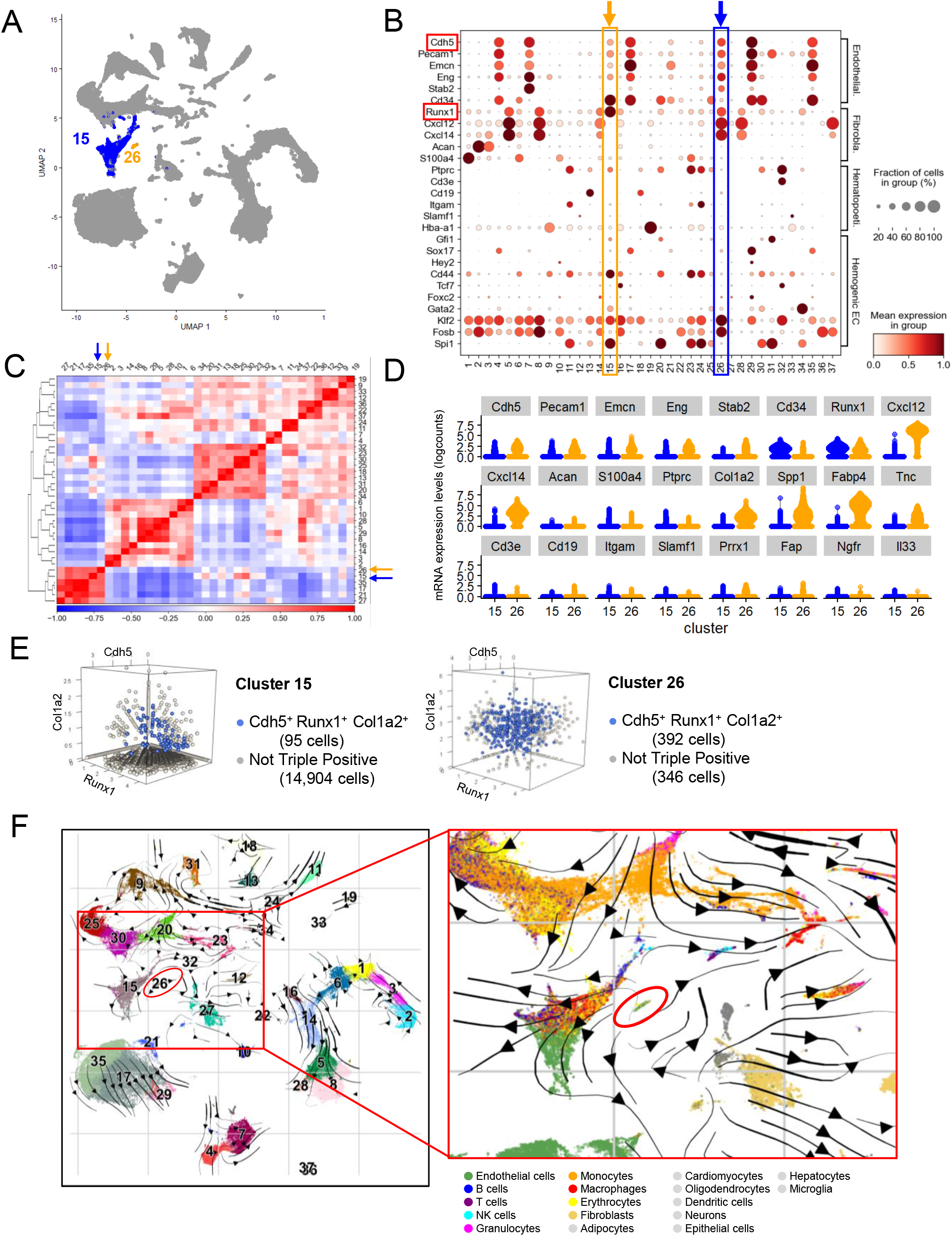
Single-cell transcriptomic analysis of prospective hemogenic endothelial cells. (A) Uniform Manifold Approximation and Projection (UMAP) representation of clusters 15 and 26 identified by unsupervised clustering. The transcriptomic data is inclusive of six data sets. (B) Expression levels of selected marker genes for endothelial cells, fibroblasts, hematopoietic cells, and hemogenic endothelium in each cell cluster. Dot plot representation of gene expression levels and fraction of expressing cells. (C) Pearson correlation matrix of all clusters, based on the top-50 principal components. (D) Selected gene expression profiles of cells in clusters 15 (blue) and 26 (orange) represented as violin plots. (E) Three-dimensional visualization of single cells co-expressing Cdh5, Runx1 and Col1a2 in clusters 15 and 26. (F) Single-cell RNA velocity plot generated by scVelo dynamical modeling showing cluster relationships projected onto UMAP. The left panel shows color-coded clusters. The right panel shows magnification and color-coded cell types. Cluster 26 is enclosed in the red oval. Arrows represent predicted trajectories. See also Figure S3.

A distinguishing feature of the “adult” cluster-26 from the “embryonic” cluster-15 was the higher expression of the mesenchymal markers Col1a2, Spp1, Fabp4 and Tnc, in addition to Cxcl12 and Cxcl14 (Figure 2B and 2D), all included among the top marker genes that distinguish the clusters (Figure S3E). Noteworthy, cluster 26 minimally expresses Ngfr or Il33 (Figure 2D), marker genes of a BM cell population that displays endothelial and stromal markers and exerts hematopoietic niche functions (Kenswil et al., 2021). Additional analyses showed that cells of cluster 26 are mostly quiescent whereas cluster 15 cells are actively proliferating (Figure S3F).

Also, whereas most (53.1%) cells of “adult” cluster-26 co-expressed Cdh5, Runx1 and Col1a2, a small minority (<1%) of “embryonic” cluster 15 co-expressed Cdh5, Runx1 and Col1a2 (Figure 2E), separating “adult” from “embryonic” Runx1-expressing ECs. Gene set enrichment analysis (GSEA) and gene ontology (GO) term analysis (Mootha et al., 2003; Subramanian et al., 2005) revealed substantial enrichment of transcriptional signatures associated with “vasculature development” and “hematopoietic progenitor cell differentiation” in cells of the adult cluster 26 (Figure S3G). Substantiating this transitional potential, RNA velocity (Bergen et al., 2020) revealed that cells of cluster 26 may have the potential to be hemogenic (Figure 2F). Thus, we will henceforth refer to cell cluster 26 as provisionally reflecting “adult hemogenic endothelial cells (AHEC)”. We examined whether a similar population exists in tissues other than the BM by analyzing public single cell data sets from 11 tissues (Kalucka et al., 2020). Only 3 cells co-expressing Cdh5, Runx1 and Col1a2 were detected among 32,000 single-cells, suggesting that AHEC are restricted to the BM (Figure S3H).

### Adult hemogenic endothelial cells are transplantable

Next, we looked for the presence of cells resembling the putative AHEC cluster 26 in the adult BM. Since bioinformatic analysis identified Col1a2 as a marker of putative AHEC, we took advantage of the inducible Col1a2 Cre-ER mouse strain (Zheng et al., 2002) to generate the mouse lines Col1a2-Cre^ERT2^/mTmG and Col1a2-Cre^ERT2^/ZsGreen for tracking cells expressing Col1a2 gene and their progeny (Figure 3A). Four weeks after tamoxifen administration (at 3-4 weeks of age), Col1a2-Cre^ERT2^/ZsGreen mice displayed a population of ZsGreen^+^ cells in the BM (Figure S4A and S4B). Among the BM ZsGreen^+^ (Col1a2) cells, we detected a distinct population of CD45^-^VE-Cadherin^+^RUNX1^+^ cells comprising 1-2% of all BM CD45^-^VE-Cadherin^+^ cells (Figure 3B). A similar population was detected in the BM of adult (age 8-12 weeks) wild-type C57Bl/6 mice (Figure 3C) and Cdh5-Cre^ERT2^(PAC)/ZsGreen mice (Figure 3D), accounting for a similar proportion (∼2%) of all BM ECs. Consistent with these results, we detected rare CD45^-^Endomucin^+^ZsGreen^+^cells in the BM of adult Col1a2-Cre^ERT2^/ZsGreen mice induced with tamoxifen (Figure 3E).

**Figure 3.**
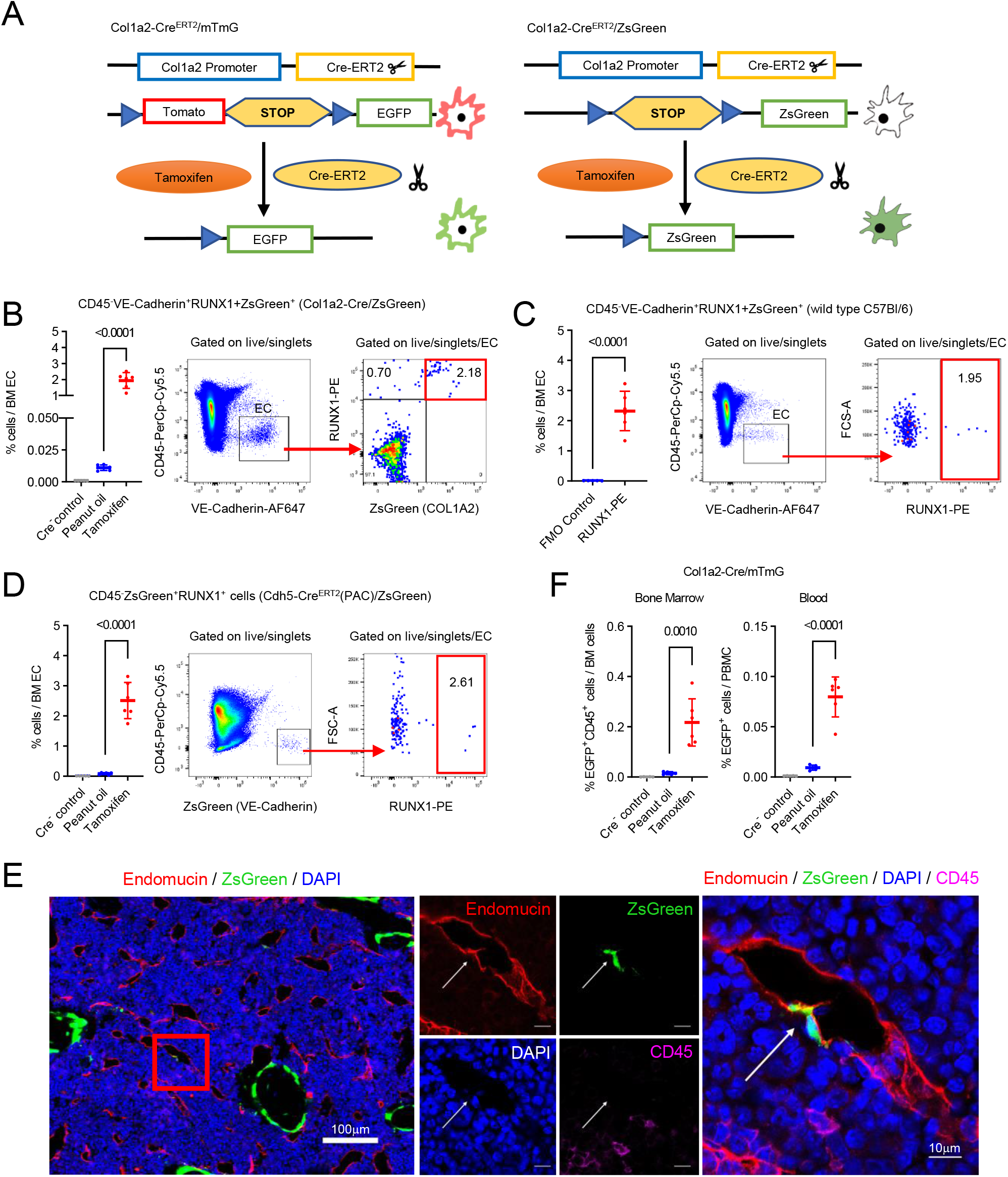
Putative hemogenic endothelial cells are detected in adult mouse bone marrow. (A) Schematic representation of the Col1a2 tracking lines. (B – D) Identification of RUNX1^+^ endothelial cells in the bone marrow of Col1a2-Cre^ERT2^/ZsGreen adult mice (B), wild type C57Bl/6 mice (C), and Cdh5-Cre^ERT2^(PAC)/ZsGreen mice (D); results shown as mean percent of all BM endothelial cells in each group. Each dot represents results from individual mice (1 femur plus 1 tibia combined). Representative gating strategies are shown in the middle and right panels. Statistical significance of group differences is indicated by the *P* value (unpaired two-tailed Student’s *t*-test). FMO: Fluorescence Minus One control. (E) Representative confocal microscopy image of bone marrow section from a Col1a2-Cre^ERT2^/ZsGreen adult mouse induced with tamoxifen showing an Endomucin^+^ZsGreen^+^ cell lining a vascular structure. (F) Percent EGFP^+^CD45^+^ cells in bone marrow and blood of Col1a2-Cre^ERT2^/mTmG control and tamoxifen treated Cre^+^ mice. The results are expressed as mean percent; each dot represents results from individual mice (1 femur plus 1 tibia combined). See also Figure S4

To investigate if this population has hemogenic potential, we looked for hematopoietic CD45^+^ cells within the Col1a2-tracked population after tamoxifen induction. By flow cytometry, we detected subsets of CD45^+^EGFP^+^ (Figure 3F) and CD45^+^ZsGreen^+^ (Figure S4C) cells in the BM and blood from Col1a2-Cre^ERT2^/mTmG and Col1a2-Cre^ERT2^/ZsGreen mice. Also, we detected subsets of CD45^+^EGFP^+^ and CD45^+^ZsGreen^+^ cells in the blood of Col1a2-Cre^ERT2^/ZsGreen and Col1a2-Cre^ERT2^/mTmG mice, supporting the hemogenic potential of Col1a2-tracked cells (Figure S4D and S4E).

To further evaluate these observations, we sorted the VE-Cadherin^+^CD45^-^ ZsGreen (Col1a2)^+^ cells from the BM of adult (age 7-9 weeks) Col1a2-Cre^ERT2^/ZsGreen mice four weeks after tamoxifen administration (Figure 4A). This sorted population of VE-Cadherin^+^CD45^-^ZsGreen (Col1a2)^+^ cells expressed Cdh5, Col1a2, Cxcl12, Runx1 and Spp1 (osteopontin) mRNAs to varying degrees, and thus differed from unselected ECs, VE-Cadherin^-^CD45^-^ “stromal” cells, and CD45^+^ hematopoietic cells (Figure 4B). Since concerted efforts to culture and immortalize these sorted VE-Cadherin^+^CD45^-^ ZsGreen (Col1a2)^+^ cells were unsuccessful (Supplemental Table 1), we transplanted these cells (10-20×10^3^ cells/mouse) into adult (8 weeks old) wild type C57Bl/6 mice (Figure 4C) and looked for their survival in the recipients. Fluorouracil (5-FU)-conditioned transplant recipients of VE-Cadherin^+^CD45^-^ ZsGreen (Col1a2)^+^ cells contained ZsGreen^+^CD45^+^ cells in BM and blood four weeks after the transplant (Figure 4D), indicating that these CD45^-^ ECs had produced CD45^+^ hematopoietic cell progeny after been transplanted. The recovery of ZsGreen^+^CD45^+^ cells in BM and blood was significantly greater in transplant recipients of VE-Cadherin^+^CD45^-^ZsGreen (Col1a2)^+^ cells compared to transplant recipients of unselected adult ECs (CD45^-^VE-Cadherin^+^ZsGreen^+^ ECs from 7-9 weeks old Cdh5-Cre^ERT2^(PAC)/ZsGreen mice) (Figure 4D), indicating enrichment of adult ECs with hemogenic potential. Phenotypically, the fluorescent CD45^+^ cells identified in the recipients of putative AHEC from the Cdh5-Cre^ERT2^(PAC)/ZsGreen) and Col1a2-Cre^ERT2^/mTmG mice were predominantly CD11b^+^ granulocytes and monocytes with very few CD19^+^ and CD3^+^ lymphocytes in BM and blood (Figure 4E and Figure S4F). Instead, the recipients of ECs without further selection (VE-Cadherin^+^CD45^-^ ZsGreen (Cdh5)^+^ cells) contained some CD19^+^ and CD3^+^ lymphocytes among the ZsGreen^+^CD45^+^ cells in the BM and blood (Figure 4E), suggesting that the Cdh5 population has a greater multilineage potential compared to the Col1a2 population. We also visualized rare CD45^+^ CD11b^+^ZsGreen^+^ cells in the BM and blood of mice transplanted with VE-Cadherin^+^CD45^-^ZsGreen (Col1a2)^+^ cells (Figure 4F and 4G).

**Figure 4.**
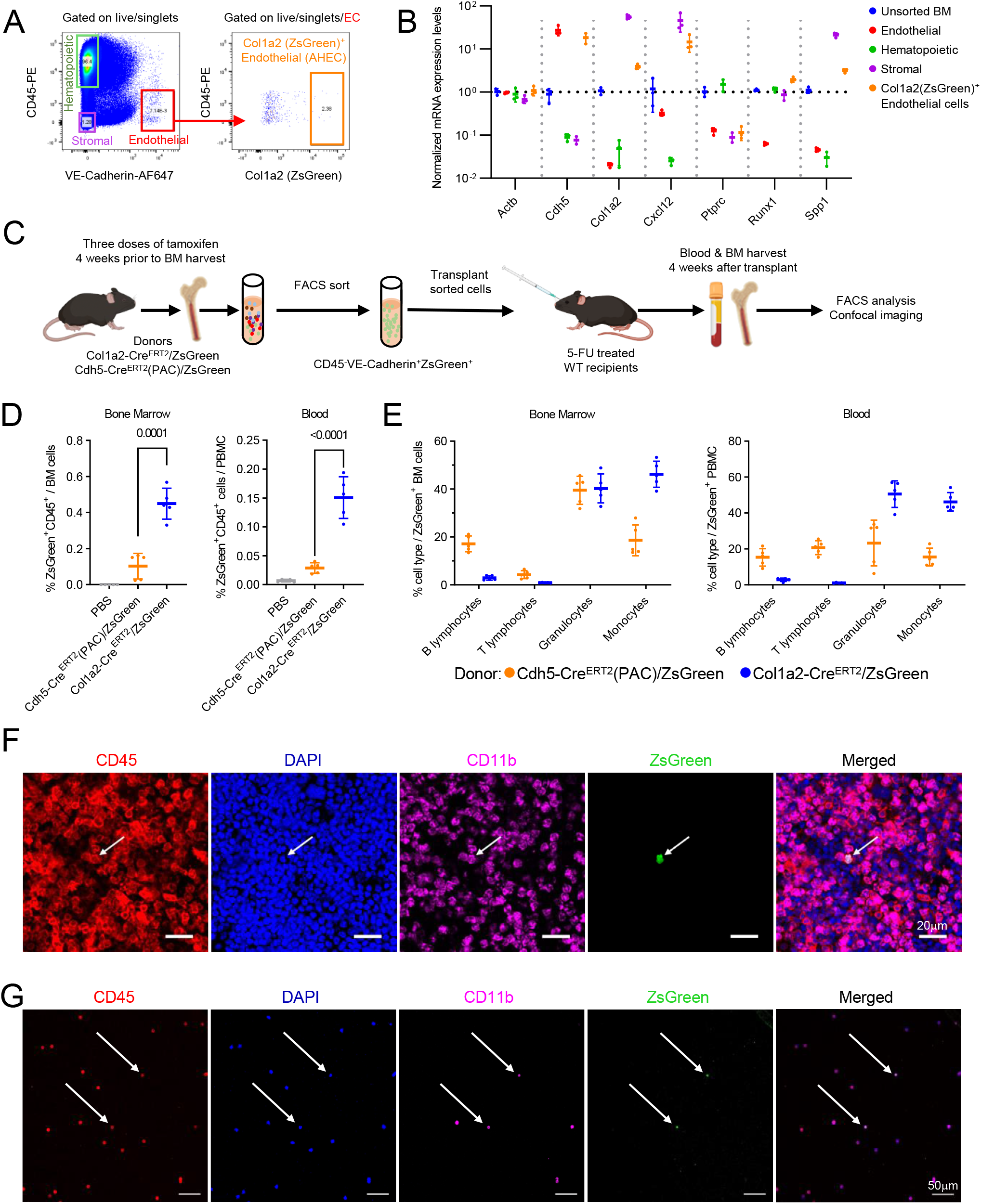
Putative adult hemogenic endothelial cells are hemogenic in transplant recipients. (A) FACS sort gating for populations of hematopoietic cells, endothelial cells, stromal cells and Col1a2-tracked endothelial cells from tamoxifen treated Col1a2-Cre^ERT2^/ZsGreen mice. Representative experiment. (B) Gene expression profiling of unsorted BM and sorted BM populations defined in (A) by qRT-PCR. Normalized by Gapdh and unsorted BM. Dots reflect experimental triplicates. (C) Schematic of experimental design for BM transplantation; donor cell procurement, sort-selection, recipients conditioning, and transplant. (D and E) Percent donor derived ZsGreen^+^CD45^+^ cells detected in BM and blood of recipients C57Bl/6 mice transplanted with no cells (PBS only), sorted CD45^-^VE-Cadherin^+^ZsGreen(Cdh5)^+^ cells (orange) or sorted bone marrow VE-Cadherin^+^CD45^-^ZSGreen (Col1a2)^+^ cells (blue). The results are expressed as percent of recipient BM cells and PBMC (D); Relative abundance of B and T lymphocytes, granulocytes, and monocytes within the ZsGreen^+^CD45^+^ cells in BM and blood of transplant recipients (E). Results of individual transplant recipients are shown as dots; group means with SD are shown; statistical significance of group differences is indicated by *P* values (unpaired two-tailed Student’s *t*-test). (F, G) Representative confocal microscopy images showing CD45^+^ZsGreen^+^CD11b^+^ cells (pointed by the arrows) in BM section (F) and blood (G) of transplant recipients of VE-Cadherin^+^CD45^-^ZsGreen (Col1a2)^+^ cells.

In sum, BM CD45^-^VE-cadherin^+^ ECs generate hematopoietic progenitor cells (LSK) and mature blood cells in wild type transplant recipients.

## DISCUSSION

Our results identify the presence of hemogenic ECs in the BM of adult mice that generate hematopoietic progenitors, mature lymphocytes, granulocytes, and monocytes. Previously, hemogenic ECs were identified during embryonic development or perinatally but not thereafter (Yvernogeau *et al*., 2019). The current finding argues that EHT is not limited to the prenatal or perinatal development but is detected at 28 weeks after birth. This adult EHT does not include a demonstrable output of HSC (CD150^+^ LSK cells), and thus differs from embryonic EHT that arises from the arterial endothelium of the aorta-gonad-mesonephros and the vitelline and umbilical arteries contributing to HSC and multipotent progenitors. Whether this restricted potential is a consequence of intrinsic characteristics of the EC population identified here, which shares mesenchymal markers and expresses Runx1, or a consequence of niche factors in the adult BM remains to be investigated.

Hematopoietic progenitors contributing to production of adult lymphoid and myeloid cells have generally been described as descendent from definitive and long-lasting HSC generated during embryonic development or perinatally. However, transient progenitors with macrophage, granulocyte and fetal erythrocyte potential emerge from the yolk sac before definitive HSC emerge (Gomez Perdiguero et al., 2015; Swiers et al., 2013) and others with preferential lymphocyte potential emerge after the emergence of definitive HSCs (Beaudin et al., 2016). Recently, a not-transplantable population of multipotent progenitors that provides most lymphoid cells throughout adult life was identified arising from embryonic arteries when definitive HSC also emerge (Patel *et al*., 2022). Our detection of transient hematopoietic progenitors originating from transplantable adult ECs identifies a previously unrecognized source of myeloid and lymphocytic cells in the blood and perhaps in other tissues. Altogether, our results reveal a functional capability of ECs in the adult BM that generates blood cells. These results suggest that hematopoiesis in the adult may arise through the contribution of cells and processes beyond the HSCs generated through aortic EHT during development.

### Limitations of the study

Our experiments show the existence of hemogenic ECs in the adult BM but have not investigated how EHT is regulated in these cells. The lifespan and function of the hematopoietic progenitors generated through adult EHT are unclear, and we could not expand adult hemogenic ECs in culture as we have no knowledge of niche factors that sustain these cells in the BM. Improved tracking systems, such as the DRE-rox alone or combined with Cre-lox, and generation of a Runx1-EGFP mouse line would be helpful to confirm and extend the current results.

## ACKNOWLEDGEMENTS

This project is supported by the Intramural Program of the Center for Cancer Research, National Cancer Institute, National Institutes of Health. This work used the computational resources of the NIH High Performance Computing (HPC) Biowulf cluster (*http://hpc.nih.gov*) and Frederick

Research Computing Environment (FRCE). Flow cytometry and cell sorting were performed at the Center for Cancer Research (CCR) Flow Cytometry Core Facility and microscopy analyses were done at the Center for Cancer Research (CCR) Microscopy Core Facility in Building 37 of the National Cancer Institute (NCI), NIH. We thank Drs. Ralf Adams and Manfred Boehm for mouse line *Cdh5-Cre*^*ERT2*^*(PAC)*; Drs. Yoshiaki Kubota and Yosuke Mukoyama for mouse line *Cdh5-Cre*^*ERT2*^*(BAC)*. We thank Dr. Subhadra Banerjee, Dr. Shafiuddin Siddiqui and Ms. Karen M. Wolcott for providing flow cytometry support and Dr. Michael Kruhlak and Mr. Langston Lim for confocal microscopy support. We thank Drs. Douglas Lowy, Michael DiPrima and Hidetaka Ohnuki for thoughtfully commenting on the experimental design and other aspects of this work.

## AUTHOR CONTRIBUTIONS

J-X F. and G. T. formulated the initial concept. J-X F. designed and executed experiments. J-X F. and G. T. analyzed the data. CC. L. and F. L. performed flow cytometry data acquisition and cell sort. J-X F., M-T Y., D. W., and A. A. performed computational analyses. G. T. and J-X F. wrote the manuscript. All authors reviewed the manuscript and provided comments.

## FIGURE TITLES AND LEGENDS

## STAR METHODS RESOURCE AVAILABILITY

### Lead contact

Further information and requests for resources and reagents should be directed to and will be fulfilled by the lead contact, Giovanna Tosato

### Materials availability

All the materials will be available upon request to the lead contact. An NCI/NIH material transfer agreement may be required. Fee(s) for shipping and/or handling might apply.

### Data and code availability

No new RNA sequencing dataset was generated by this study. The published single-cell RNA sequencing datasets used in this research are: GSE108885, GSE108891, GSE118436, GSE123078, GSE122465, GSE128423, GSE145477, GSE156635, GSE137116, E-MTAB-8077.

No custom software was developed for this study. Methods describe the used analytical tools and variables. Bash, R, and Python codes are available from the lead contact upon reasonable request.

## EXPERIMENTAL MODEL AND SUBJECT DETAILS

### Mouse strains

All animal studies were approved by the Institutional Animal Care and Use Committee of the CCR (Bethesda, MD), National Cancer Institute (NCI), NIH and conducted in adherence to the NIH Guide for the Care and Use of Laboratory Animals (National Academies Press, 2011) and approved protocols. *Cdh5-Cre*^*ERT2*^*(PAC)* mice (Sorensen *et al*., 2009) (MGI:3848982, PAC: P1 artificial chromosome) were a gift of Dr. R. Adams and *Cdh5-Cre*^*ERT2*^*(BAC)* mice (Okabe *et al*., 2014) (MGI:5705396, BAC: bacterial artificial chromosome) were a gift of Dr. Kubota. *Col1a2-CreER* mice (Zheng *et al*., 2002) were purchased from the Jackson Laboratory (JAX#029567). Cre-dependent *Ai6-ZsGreen* (Madisen *et al*., 2010) (JAX#007906) and *mTmG* (Muzumdar *et al*., 2007) (JAX#007676) fluorescent reporter mice were purchased from the Jackson Laboratory. In *ZsGreen* reporter mice, Cre activity leads to constitutive expression of ZsGreen1 in cell bodies. In *mTmG* reporter mice, Cre activity leads to an irreversible switch from cell membrane-localized tdTomato(mT) to membrane-localized EGFP (mG).

All animals were bred in the animal facilities of CCR/NCI (Bethesda, MD), where rodent pathogens are quarterly tested. The mice were maintained in a predominantly C57Bl/6J background. Mice were identified with ear tags (Braintree Scientific, #EP10051) and routinely genotyped by PCR. Protocols and primer sequences can be found in the related references and will be provided upon request. No mouse was excluded from the experiments, unless assessed sick by the veterinarians or fight wounds were found at harvest. Tamoxifen (Sigma-Aldrich, #T5648) dissolved in peanut oil (Sigma-Aldrich, #P2144) (10mg·mL^-1^) was administered orally (via gavage using 22g feeding needles) at 100mg·kg^-1^. Unless otherwise specified, three doses in consecutive days were administered. Unless indicated otherwise, 4-to 6-week-old male and female mice were used. Littermates of the same sex were randomly assigned to experimental groups. Mice were typically sacrificed between 9LAM and 11LAM local time.

### Transplantation experiments

Four days prior to transplant, recipient mice received one dose of 5-FU (150 mg·kg^-1^ in PBS, Sigma-Aldrich, #F6627) intraperitoneally under isoflurane anesthesia. Sorted bone marrow cells (5,000 to 20,000 cells in 100μL PBS) were inoculated retro-orbitally under isoflurane anesthesia. Bone marrows and blood were harvested from transplant recipients four weeks after the transplant unless otherwise specified.

## METHOD DETAILS

### Blood collection, counts, and analysis

Blood was collected from the mouse abdominal aorta with BD Vacutainer™ EDTA tubes (BD #367856) and Vaculet™ blood collection needles (23G, EXELINT #26766). Blood smears were prepared with 10 μL collected blood. For flow cytometry analysis, ACK buffer (Lonza, #BP10-548E) was added to the blood to lyse red blood cells before Fc receptor blocking and antibody staining. For non-terminal blood collection, ∼20 -50μL blood was collected by submandibular blood sampling, using a 3mm animal lancet (BRAINTREE SCI., GR-3MM) and a 250μL BD Microtainer^®^ K2EDTA tube (BD #365974). Kwik Stop^®^ Styptic Powder was applied to stop the bleeding (Miracle Corp. #423615).

### Bone marrow harvesting

Bone marrow was harvested using one of the two methods described below. For hematopoietic cells, bone marrows were harvested by flushing mouse femurs and tibiae with ice-cold Sort Buffer (5mM EDTA, 25mM HEPES, 2% fetal bovine serum (FBS, SIGMA, #F2442) in 1× PBS (GIBCO, #10010-031)), followed by red cell lysis with ACK buffer. Cells were then washed with Sort Buffer and passed through a 40μm cell strainer (GREINER BIO-ONE, #542040, #542140). For greater preservation of endothelial cells, bone marrows were harvested by gently crushing mouse femurs and tibiae in Sort Buffer (5mM EDTA, 25mM HEPES, 2% FBS in 1× PBS). Red cells lysis was performed using ACK buffer. Bone marrow cells were then incubated with 0.1U·mL^-1^ Collagenase (Worthington Biomedical Corp., #LS004176), 0.8U·mL^-1^ Dispase (Worthington Biomedical Corp., #LS02109), and 0.5mg·mL^-1^ DNase (Worthington Biomedical Corp., #LS006344) in 1x Hanks’ Balanced Salt Solution (HBSS) with Ca^2+^ and Mg^2+^ (Gibco, # 14065056) at 37°C for 30 min on a rotating mixer. Cells were then washed with Sort Buffer and passed through a 40μm cell strainer.

### Flow cytometry and cell sorting

For intracellular antigen detection, single cell suspensions of bone marrow and blood were first incubated with Azide-Free Fc Receptor Blocker (Innovex, #NB335-60), following the manufacturer’s instructions. After washing, cells were first stained with surface marker antibodies (listed in Supplemental Table 3) at the concentration of 2μg per 1×10^7^ cells in Sort Buffer for 30 minutes at 4°C, and then stained with live/dead cell discriminating BioLegend Zombie Dyes (UV, NIR, Violet, or Yellow, BioLegend #423108, #423106, #423114, and # 423104) following the manufacturer’s instructions. After washing, cells were fixed in 4% paraformaldehyde for 10 minutes at 37°C, and then permeabilized with 1% saponin (Sigma-Aldrich, # 47036) /Sort Buffer for 30min on ice. Subsequently, the cells were stained with rat monoclonal primary RUNX1-PE Ab (Invitrogen, # 12-9816-80) in 0.1% saponin/Sort Buffer overnight. After washing and resuspension, propidium iodide (PI, 0.5μM, Millipore Sigma, # P4170), 7-AAD (1μg·mL^-1^, Millipore Sigma, #A9400) or DAPI (0.5μg·mL^-1^, BioLegend, #422801) was added as a nuclear counterstain. For live cell staining without cell permeabilization, after cell surface antibody staining, cells were washed and suspended in Sort buffer containing propidium iodide (PI, 0.5μM, Millipore Sigma, # P4170), 7-AAD (1μg·mL^-1^, Millipore Sigma, #A9400) or DAPI (0.5μg·mL^-1^, BioLegend, # 422801), to distinguish dead cells from the live cells. Compensation beads (BD Biosciences, #552844) were used for flow cytometer compensation. Flow cytometric data were acquired with BD FACSCanto-II, BD LSRFortessa, BD FACSymphony A5 (BD Biosciences), Sony SA3800 or Sony ID7000 cell analyzers. Cell sorting was performed with BD FACS Aria III, BD FACS Aria Fusion or Sony SH800S cell sorters. FSC and SSC profiles were used for excluding dead cells and debris. 7-AAD, PI, DAPI or BioLegend Zombie Dye was used for excluding dead cells. FSC-W versus FSC-H and SSC-W versus SSC-H were used to gate on single cells. Data were analyzed with FlowJo (BD, v10.8.1), SONY ID7000 Software (Version 1.2.0.28212) or FACS Diva (BD, v6.1 and v9.0).

For FACS sorting of endothelial cells, after Fc receptor blocking, bone marrow cells were first depleted of CD45^+^ cells with MojoSort mouse CD45 nanobeads (BioLegend #480028) following the manufacturer’s protocol, and then stained with specific antibodies.

### Bone marrow cryosections

Deeply anesthetized mice were transcardially perfused with 20mL ice-cold 1x PBS, followed by perfusion with 15mL ice-cold hydrogel solution (5% acrylamide/bis-acrylamide 19:1 (Sigma-Aldrich, #A2917), 2.5mg/mL polymerization initiator VA-044 (FUJIFILM, Wako, VA-044, Water soluble Azo initiators), 4% PFA in 1× PBS). Femurs and tibiae were collected in tubes containing 5mL hydrogel solution and incubated at 4°C for 4 hours. The bones were then washed with PBS and incubated at 37°C for 2 hours. Bone decalcification was performed by incubating the bones in 40mL 0.5M EDTA pH 8.0 (KD Medical, #RGF-3130) for 3 days on a rotate mixer, with daily refreshed 0.5M EDTA solution. The bones were then dehydrated in 20% sucrose and 2% polyvinylpyrrolidone in PBS overnight. Dehydrated bones were then embedded in OCT (SAKURA, #4583) blocks using Precision Cryoembedding System (IHC WORLD, #IW-P101).

Cryosections (10μm) for immunofluorescence staining were prepared from OCT frozen bone blocks using Leica CM3050S microtome, low-profile microtome blades (Leica 819, #14035838925), and TruBond™ 380 adhesion slides (Electron Microscopy Sciences, # 63701-W10).

### Immunofluorescence staining and imaging

Tissue sections were rehydrated with 1× PBS (15 minutes), permeabilized in 0.3% Triton X-100 (Sigma-Aldrich, #T9284)/PBS (15 minutes), washed in 1× PBS, and incubated (2 hours) with blocking solution (2% BSA, 5% donkey serum (SIGMA, #D9663), and 0.3% Triton X-100/PBS). Samples were then washed 3 times with PBS and incubated with primary antibodies (5ng/mL; 4°C overnight). When secondary antibodies were used, 3 PBS washes were performed before incubating with fluorescent-labeled secondary antibodies (2ng/mL, room temperature, 2 hours). After washing (3x, 10 minutes each with 1× PBS), DAPI was added (300nM in PBS, 10 min). After 3 washes (5 min each with 1× PBS), coverslips were mounted (EPREDIA, #9990402), dried and sealed with nail polish. For blood smear staining, slides were first soaked in acetone/methanol/PFA (19:19:2 for 90 seconds) before rehydration(Happerfield et al., 2008). Confocal imaging was performed with Zeiss LSM 780, Zeiss LSM 880 NLO Two Photon, or Nikon *ECLIPSE* Ti2-E SoRa systems. Images were processed with Zen (Zen Black v2.3, release Version 14.0.12.201, Zen Blue Lite v2.5, Carl Zeiss), Bitplane Imaris (v9.7.0, Oxford instruments), Microsoft 365 PowerPoint (Version 2206, Build 15330.20264, for whole image contrast and brightness adjustment, cropping and aligning) and Photoshop (v23.3.0, Adobe, for whole image contrast and brightness adjustments).

### Single cell RNA-seq data analysis

Raw FASTQ and BAM files were downloaded from publicly available datasets (GSE108885(Tikhonova *et al*., 2019), GSE108891(Tikhonova *et al*., 2019), GSE118436(Tikhonova *et al*., 2019), GSE123078(Tikhonova *et al*., 2019), GSE122465(Baccin *et al*., 2020), GSE128423(Baryawno *et al*., 2019), GSE145477(Kalucka *et al*., 2020), GSE156635(Sivaraj *et al*., 2021), GSE137116(Zhu *et al*., 2020), E-MTAB-8077 (Kalucka *et al*., 2020)). The BAM files were first converted to FASTQ files with bamtofastq (10x Genomics, v1.4.1). All FASTQ files were then processed by the count function of Cell Ranger (10x Genomics, v6.1.2) and aligned to the mouse genome (mm10, version 2020-A), to generate read matrixes. Quality control was performed based on mitochondrial reads percentage and gene numbers contained using scater package (v1.22.0(McCarthy et al., 2017)) with quickPerCellQC function. Normalization was done with scuttle::logNormCounts (1.4.0(McCarthy *et al*., 2017)), using the predetermined size.factor (SingleCellExperiment::sizeFactors, 1.18.0(Amezquita et al., 2020)). Batch effect was corrected with batchelor package (1.10.0(Haghverdi et al., 2018)) using FastMnn algorithm (k=20). Unsupervised clustering was done using Louvain method of igraph R package (v1.3.2(Csardi and Nepusz, 2006), resolution = 1), using batch corrected values from Cell Ranger. UMAP coordinates were determined using normalized batch corrected expression values from Cell Ranger (scatter, v1.22.0(McCarthy *et al*., 2017)). Cell cycle analysis was performed with SingleR (v1.10.0(Aran et al., 2019)) using cell cycle Gene Ontology dataset (org.Mm.eg.db, v3.15.0 (Carlson et al., 2019), GO:0007049) and scRNAseq(v2.10.0 (Risso et al., 2021))::BuettnerESCData()(Buettner et al., 2015) and as references. Marker genes were identified using scanpy (v1.9.1(Wolf et al., 2018)) with t-test method. Cell type was annotated with SingleR (v1.10.0) using MouseRNAseqData(Benayoun et al., 2019) from celldex package

(v1.4.0(Aran *et al*., 2019)). GSEA was performed with presto (v1.0.0(Korsunsky et al., 2019)) and fgsea (v1.20.0(Korotkevich et al., 2021)). Trajectory analysis was performed using scanpy.tl.paga(Wolf et al., 2019) from scanpy package. For RNA velocity assay, Cell Ranger processed bam files were first sorted with samtools (v1.15.1(Danecek et al., 2021)), then processed with velocyto (v0.17(La Manno et al., 2018)) and mouse genome (mm10, version 2020-A) to generate loom files containing spliced and unspliced reads. Loom files were then processed with scvelo (v0.2.4(Bergen *et al*., 2020)) using its dynamical modeling (pp.filter_and_normalize: min_shared_counts=20, n_top_genes=2000, pp.moments: n_pcs=30, n_neighbors=30). Data visualization was plotted with scater, seurat, scanpy, scvelo, ggplot2 (v3.3.6(Wickham, 2016)), Matplotlib (v3.5.2(Price-Whelan et al., 2018)), and rgl (v0.109.2(Adler et al., 2003)). This work utilized the computational resources of the NIH HPC Biowulf cluster (*http://hpc.nih.gov*) and Frederick Research Computing Environment (FRCE). Some local computing was done with macOS 12 Monterey (v12.5, Build 21G72), Ubuntu (22.04 LTS), Windows 10/11 WSL2 (Ubuntu 20.04 LTS). Python version: 3.8. R version 4.2.

### RNA isolation and quantitative RT PCR

RNA was extracted using RNeasy^®^ Micro Kit (QIAGEN, #74004), following the manufacturer’s protocol. Cells were sorted directly into RNA lysis buffer (Buffer RTL of RNeasy^®^ Micro Kit). cDNA samples were prepared with SuperScript IV Reverse Transcriptase (Invitrogen, #18091050), following manufacturer’s instructions. Real Time PCR was performed using Applied Biosystems™ TaqMan™ Fast Advanced Master Mix (4444557) and Applied Biosystem™ QuantStudio™ 5 Real-Time PCR System. TaqMan™ probes used were purchased from Applied Biosystems™: Spp1 (Mm00436767_m1); Cxcl12 (Mm00445553_m1); Col1a2 (Mm00483888_m1); Cdh5 (Mm00486938_m1); Runx1 (Mm01213404_m1); Ptprc (Mm01293577_m1); Gapdh (Mm99999915_g1); Actb (Mm02619580_g1). The reaction condition was set as follows: 50°C 2 minutes, 95°C 20 seconds, 45 cycles of 95°C 1 second, 60°C 20 seconds. Ct values were determined using the ABI QuantStudio™ Design & Analysis Software (v1.5.2). Relative gene expression was assessed using the 2^-ΔΔCt^ method, normalized to Gapdh expression level for each sample. The data was further normalized to gene expression levels in the unsorted bone marrow sample to calculate relative gene expression levels in each sample. Data reflect triplicates real time PRC experiments.

## QUANTIFICATION AND STATISTICAL ANALYSIS

No statistical method was used to predetermine sample size. No data were excluded from the analyses. Mice with the correct genotypes were randomly assigned to control or treated groups. All data are represented as meanL±LS.D. Comparisons between two groups were performed using Student’s paired or unpaired *t*-tests. Statistical analyses were performed with GraphPad Prism (v9.0.1). A statistical difference of *P*<0.05 was considered significant.

## KEY RESOURCES TABLE

Uploaded separately.

## SUPPLEMENTAL FIGURE TITLES AND LEGENDS

**Figure S1 related to Figure 1. Endothelial cells contribution to hematopoiesis in adult bone marrow is revealed by Cdh5-Cre**^**ERT2**^ **mouse tracking lines**.

(A) Bone marrow (BM) flow cytometry analysis of adult Cdh5-Cre ^ERT2^(PAC) /mTmG mice demonstrates that EGFP fluorescence identifies most CD45^-^VE-Cadherin^+^CD31^+^ endothelial cells (EC) four weeks after tamoxifen administration (left). Each data-point represents results from individual mice (1 femur plus 1 tibia combined). The gating strategy for the flow cytometry analysis is shown in a representative example from a tamoxifen-treated mouse (mid and right panels). EC were first gated (live/singlets/CD45^-^ VE-cadherin^+^) and the proportion of EGFP^+^CD31^+^ cells within the EC gate was measured.

(B) Flow cytometry analysis of adult BM from Cdh5-Cre^ERT2^(PAC)/ZsGreen and Cdh5-Cre^ERT2^(BAC)/ZsGreen mice shows that ZsGreen fluorescence identifies most EC four weeks after tamoxifen administration, but also tracks a small proportion of EC expressing tamoxifen-independent fluorescence in Cre^+^ but not Cre^-^ mice. Each data-point represents results from individual mice (1 femur plus 1 tibia combined). The gating strategy for the flow cytometry analysis is shown in a representative example from a tamoxifen-treated mouse (middle and right panels).

(C) Confocal microscopy images of a representative BM section from a Cdh5-Cre^ERT2^(PAC)/ZsGreen adult mouse showing tamoxifen-induced ZsGreen fluorescence co-staining of most Endomucin^+^ cells. Control BM sections from a representative Cre^+^ mouse treated with peanut oil (no tamoxifen) displays occasional ZsGreen^+^Endomucin^+^ cells but no tamoxifen-independent ZsGreen fluorescence is detected in a representative Cre^-^ mouse.

(D) Confocal microscopy images of a representative BM section from a Cdh5-Cre^ERT2^(BAC)/ZsGreen adult mouse showing tamoxifen-induced ZsGreen fluorescence co-staining of most Endomucin^+^ cells. Control BM sections from a representative Cre^+^ mouse treated with peanut oil (no tamoxifen) displays occasional ZsGreen^+^Endomucin^+^ cells but no tamoxifen-independent ZsGreen fluorescence is detected in a representative Cre^-^ mouse.

(E) Percent ZsGreen^+^CD45^+^ cells within viable BM cells from Cre^-^ control, Cre^+^ control (peanut-oil treated, no tamoxifen) and Cre^+^ tamoxifen-treated mice. Each dot represents results from individual mice (1 femur plus 1 tibia combined). Each dot reflects results from individual mice (1 femur plus 1 tibia combined); the group means ± SD (error bars) are shown. *P* values (unpaired two-tailed Student’s *t*-test) are shown.

(F) Representative gating strategy for quantification of CD45^+^EGFP^+^ cells in the bone marrow of Cdh5-Cre^ERT2^(PAC)/mTmG mice treated with peanut oil only or with tamoxifen. Cre^-^ mice treated with tamoxifen were tested as controls.

(G) Representative gating strategy for quantification of CD45^+^ZsGreen^+^ cells in mouse bone marrow from Cdh5-Cre^ERT2^(BAC)/ZsGreen mice treated with peanut oil only or with tamoxifen. Cre^-^ mice treated with tamoxifen were tested as controls.

**Figure S2 related to Figure 1. Characterization of tracked hematopoietic progenitors and mature cells in adult bone marrow and peripheral blood of Cdh5-Cre reporter mice**.

(A) Percent ZsGreen^+^ cells in the bone marrow (BM) of Cdh5-Cre^ERT2^(PAC)/ZsGreen mice treated or not treated with tamoxifen showing tamoxifen-independent ZsGreen fluorescence and the significant increase of ZsGreen fluorescence induced by tamoxifen. Cre^-^ mice treated with tamoxifen were used as controls (left). Distribution of viable BM ZsGreen^+^ cells within LSK (middle). Percent B lymphocytes, T lymphocytes, granulocytes, and monocytes in all ZsGreen^+^ cells (right). Each dot reflects results from individual mice (1 femur plus 1 tibia combined); the group means ± SD (error bars) are shown. *P* values (unpaired two-tailed Student’s *t*-test) are shown.

(B) Left panel: percent ZsGreen^+^ cells in the bone marrow of Cdh5-Cre^ERT2^(PAC)/ZsGreen mice treated or not treated with tamoxifen. All other panels: the overall (ZsGreen^+^ and ZsGreen^-^) percent LSK cells, B lymphocytes, T lymphocytes, granulocytes and monocytes is similar in the bone marrow of control Cre^+^ mice (peanut oil treated, no tamoxifen) and tamoxifen treated Cre^+^ mice. This indicates that tamoxifen administration has not altered the distribution of bone marrow cell subsets analyzed. Results from individual mice are displayed as dots; means (± SD) are displayed as horizontal lines and error bars. Statistical significance of differences determined by unpaired two-tailed Student’s *t*-test are shown; *P*<0.05 is considered statistically significant; ns: not significant difference.

(C) Left panel: percent EGFP^+^ cells in the bone marrow of Cdh5-Cre^ERT2^(PAC)/mTmG mice four weeks after administration of tamoxifen or peanut oil only. Cre^-^ mice treated with tamoxifen were used as controls. All other panels: the overall (EGFP^+^ and EGFP^-^) percent LSK (Lin^-^, Sca1^+^cKit^+^), B (CD19^+^) and T (CD3^+^) lymphocytes, granulocytes (CD19^-^, CD3^-^, Ly6G^+^, CD11b^+^), and monocytes (CD19^-^, CD3^-^, Ly6G^-^, CD11b^+^) is similar in the bone marrow of control Cre^+^ mice (peanut oil treated, no tamoxifen) and tamoxifen treated Cre^+^ mice. Results from individual mice are displayed as dots; means (± SD) are displayed as horizontal lines and error bars. Statistical significance of differences determined by unpaired two-tailed Student’s *t*-test are shown; ns: not significant difference.

(D) Representative immunofluorescence image of peripheral blood ZsGreen^+^CD45^+^ nucleated (DAPI^+^) cells (pointed by the arrows) from a tamoxifen treated Cdh5-Cre^ERT2^(PAC)/ZsGreen mouse.

(F) Percent ZsGreen^+^ cells within PBMC of Cdh5-Cre^ERT2^(PAC)/ZsGreen^+^ mice treated with tamoxifen, peanut oil (no tamoxifen) or Cre^-^ mice treated with tamoxifen (left), and relative contribution of B lymphocytes, T lymphocytes, granulocytes, and monocytes to ZsGreen^+^ PBMC (right).

(F) Left panel: percent EGFP^+^ cells within peripheral blood mononuclear cells (PBMC) of Cdh5-Cre^ERT2^(PAC)/mTmG mice treated with tamoxifen or with peanut oil (no tamoxifen). Cre^-^ mice treated with tamoxifen are included as controls (left). All other panels: similar percent B lymphocytes (CD19^+^), T lymphocytes (CD3^+^), granulocytes (CD19^-^, CD3^-^, Ly6G^+^, CD11b^+^), and monocytes (CD19^-^CD3^-^Ly6G^-^CD11b^+^) is detected in PBMC of Cdh5-Cre^ERT2^(PAC)/mTmG mice treated with peanut oil only (no tamoxifen) or with tamoxifen, showing that tamoxifen administration has not altered the distribution of PBMC subsets analyzed.

(G) Similar percent B lymphocytes (CD19^+^), T lymphocytes (CD3^+^), granulocytes (CD19^-^CD3^-^ Ly6G^+^CD11b^+^), and monocytes (CD19^-^CD3^-^Ly6G^-^CD11b^+^) is measured in PBMC of Cdh5-Cre^ERT2^(PAC)/ZsGreen mice treated with peanut oil only (no tamoxifen) or with tamoxifen, showing that tamoxifen administration has not altered the distribution of PMBC subsets analyzed.

**Figure S3 related to Figure 2. Bioinformatic identification and characterization of prospective Runx1**^**+**^ **hemogenic endothelial cells in adult mouse bone marrow**.

(A) Identification of cell clusters 26 and 15 by Uniform Manifold Approximation and Projection (UMAP) visualization; the results reflect color coded unsupervised clustering of six datasets by the Louvain method.

(B) Illustration of the contribution of each of the six datasets to the overall cluster identification. The source of each dataset is identified by color and the relative contribution of each dataset to the overall cell clustering is shown.

(C) Cell type annotation is color-coded in each dataset.

(D) Data source and source contribution to all endothelial cells, adult endothelial cells, and endothelial clusters 15 and 26.

(E) Heatmap depiction of marker genes (top 15) most discriminating each cluster showing that cells in cluster 26 co-express endothelial and mesenchymal markers and the transcription factor Runx1. The orange arrow points to cluster 26; the blue arrow points to cluster 15.

(F) Cell cycle distribution of cells within each cluster showing that cluster 26 cells mostly reside in G1 whereas cells of cluster 15 mostly reside in G2M and S. Cell cycle distribution was assessed by SingleR package as detailed in the methods section.

(G) Gene set enrichment analysis (GSEA) shows that cells in cluster 26 display a selective enrichment in expression of gene sets defining Vascular Development and Hematopoietic Progenitor Cell Differentiation compared to the other clusters as reflected by the normalized enrichment score.

(H) Dataset source: single cell transcriptome atlas of murine endothelial cells from the 11 indicated tissues. Identification of cell clusters by t-distributed stochastic neighbor embedding (t-SNE) visualization. The results are shown as color-coded representation of tissue derivation (top left) and expression levels of Cdh5, Runx1 and Col1a2. The Venn diagram shows that co-expression of Cdh5, Col1a2 and Runx1 is rare in the dataset.

**Figure S4 related to Figure 3. Characterization of Col1a2-tracked cell populations in bone marrow and blood**.

(A) Representative confocal image of bone marrow from a Col1a2-Cre/ZsGreen mouse induced with tamoxifen showing that the ZsGreen^+^ cells are widely detected.

(B) In the bone marrow of Col1a2-Cre/ZsGreen mice, only few ZsGreen^+^ cells are detected in the bone marrow of a peanut oil-treated mouse (middle panels), whereas numerous ZsGreen^+^ cells are detected in the tamoxifen treated mouse (right panel). No ZsGreen^+^ cells are detected in a Cre^-^ control bone marrow (left panels). Representative images.

(C) Percent ZsGreen^+^ cells among bone marrow CD45^+^ cells (left) and peripheral blood mononuclear cells (PBMC, right) detected in Col1a2-Cre^ERT2^/ZsGreen mice after peanut oil or tamoxifen treatment. Each dot represents results from individual mice (1 femur plus 1 tibia combined). The means (± SD) are displayed as horizontal lines and error bars. Statistical significance of differences determined by unpaired two-tailed Student’s *t*-test are shown.

(D) Representative confocal image of a blood smear from a Col1a2-Cre/ZsGreen mouse treated with tamoxifen showing the presence of nucleated CD45^+^ cells tracked by ZsGreen/Col1a2 fluorescence (pointed by the arrows).

(E) Representative confocal image of a blood smear from a Col1a2-Cre/mTmG mouse treated with tamoxifen showing the presence of nucleated CD45^+^ cells tracked by EGFP/Col1a2 fluorescence (pointed by the arrows).

(F) Distribution of B and T lymphocytes, granulocytes, and monocytes in EGFP^+^ bone marrow (BM) and EGFP^+^ peripheral blood mononuclear cells (PBMC) from Col1a2-Cre/mTmG mice treated with tamoxifen. Each dot represents results from individual mice (1 femur plus 1 tibia combined).

**Supplemental Table 1, related to Figure 4. Methods to Culture and/or Immortalize AHEC**.

