## Supplemental Figures for "Bone marrow hemogenic endothelial cells contribute multilineage hematopoietic progenitors in adult mice"

### Figure S1

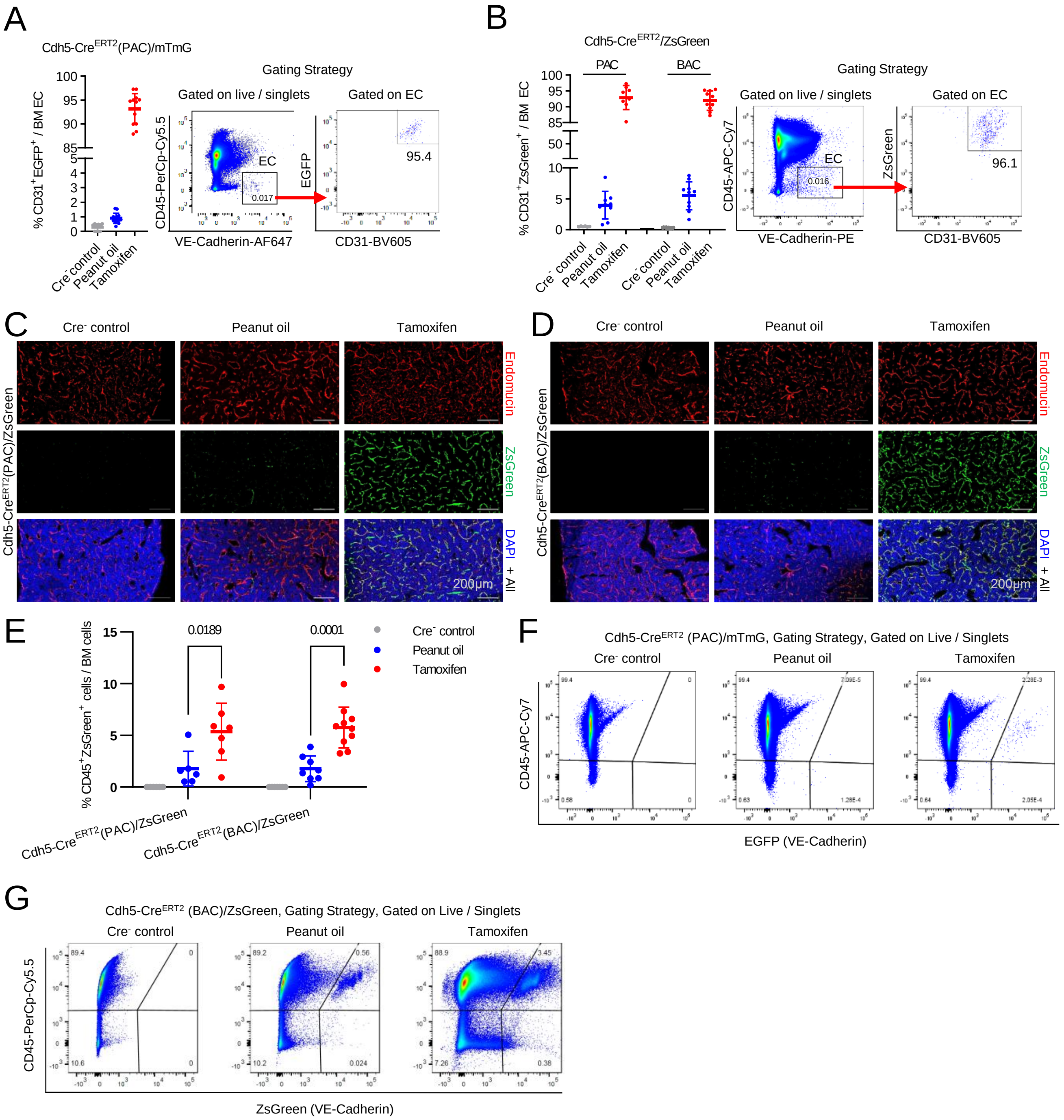

A
Cdh5-CreERT2(PAC)/mTmG
% CD31+EGFP+ / BM EC
Cre- control
Peanut oil
Tamoxifen
Gating Strategy
Gated on live / singlets
Gated on EC
CD45-PerCp-Cy5.5
VE-Cadherin-AF647
EC
0.017
EGFP
CD31-BV605
95.4
B
Cdh5-CreERT2/ZsGreen
PAC
BAC
% CD31+ZsGreen+ / BM EC
Cre- control
Peanut oil
Tamoxifen
Gating Strategy
Gated on live / singlets
Gated on EC
CD45-APC-Cy7
VE-Cadherin-PE
EC
0.016
ZsGreen
CD31-BV605
96.1
C
Cre- control
Peanut oil
Tamoxifen
Cdh5-CreERT2(PAC)/ZsGreen
Endomucin
ZsGreen
DAPI + All
200µm
D
Cre- control
Peanut oil
Tamoxifen
Cdh5-CreERT2(BAC)/ZsGreen
Endomucin
ZsGreen
DAPI + All
200µm
E
% CD45+ZsGreen+ cells / BM cells
0.0189
0.0001
Cre- control
Peanut oil
Tamoxifen
Cdh5-CreERT2(PAC)/ZsGreen
Cdh5-CreERT2(BAC)/ZsGreen
F
Cdh5-CreERT2 (PAC)/mTmG, Gating Strategy, Gated on Live / Singlets
Cre- control
Peanut oil
Tamoxifen
CD45-APC-Cy7
EGFP (VE-Cadherin)
G
Cdh5-CreERT2 (BAC)/ZsGreen, Gating Strategy, Gated on Live / Singlets
Cre- control
Peanut oil
Tamoxifen
CD45-PerCp-Cy5.5
ZsGreen (VE-Cadherin)

**Figure S1 related to Figure 1. Endothelial cells contribution to hematopoiesis in adult bone marrow is revealed by Cdh5-Cre$^{ERT2}$ mouse tracking lines.**

(A) Bone marrow (BM) flow cytometry analysis of adult Cdh5-Cre $^{ERT2}$(PAC) /mTmG mice demonstrates that EGFP fluorescence identifies most CD45$^{-}$VE-Cadherin$^{+}$CD31$^{+}$ endothelial cells (EC) four weeks after tamoxifen administration (left). Each data-point represents results from individual mice (1 femur plus 1 tibia combined). The gating strategy for the flow cytometry analysis is shown in a representative example from a tamoxifen-treated mouse (mid and right panels). ECs were first gated (live/singlets/CD45$^{-}$ VE-cadherin$^{+}$) and the proportion of EGFP$^{+}$CD31$^{+}$ cells within the EC gate was measured.

(B) Flow cytometry analysis of adult BM from Cdh5-Cre$^{ERT2}$(PAC)/ZsGreen and Cdh5-Cre$^{ERT2}$(BAC)/ZsGreen mice shows that ZsGreen fluorescence identifies most EC four weeks after tamoxifen administration, but also tracks a small proportion of EC expressing tamoxifen-independent fluorescence in Cre$^{+}$ but not in Cre$^{-}$ mice. Each data-point represents results from individual mice (1 femur plus 1 tibia combined). The gating strategy for the flow cytometry analysis is shown in a representative example from a tamoxifen-treated mouse (middle and right panels).

(C) Confocal microscopy images of a representative BM section from a Cdh5-Cre$^{ERT2}$(PAC)/ZsGreen adult mouse showing tamoxifen-induced ZsGreen fluorescence co-staining of most Endomucin$^{+}$ cells. Control BM sections from a representative Cre$^{+}$ mouse treated with peanut oil (no tamoxifen) displays occasional ZsGreen$^{+}$Endomucin$^{+}$ cells but no tamoxifen-independent ZsGreen fluorescence is detected in a representative Cre$^{-}$ mouse.

(D) Confocal microscopy images of a representative BM section from a Cdh5-Cre$^{ERT2}$(BAC)/ZsGreen adult mouse showing tamoxifen-induced ZsGreen fluorescence co-staining of most Endomucin$^{+}$ cells. Control BM sections from a representative Cre$^{+}$ mouse treated with peanut oil (no tamoxifen) displays occasional ZsGreen$^{+}$Endomucin$^{+}$ cells but no tamoxifen-independent ZsGreen fluorescence is detected in a representative Cre$^{-}$ mouse.

(E) Percent ZsGreen$^{+}$CD45$^{+}$ cells within viable BM cells from Cre$^{-}$ control, Cre$^{+}$ control (peanut-oil treated, no tamoxifen) and Cre$^{+}$ tamoxifen-treated mice. Each dot represents results from individual mice (1 femur plus 1 tibia combined). Each dot reflects results from individual mice (1 femur plus 1 tibia combined); the group means ± SD (error bars) are shown. *P* values (unpaired two-tailed Student's *t*-test) are shown.

(F) Representative gating strategy for quantification of CD45$^{+}$EGFP$^{+}$ cells in the bone marrow of Cdh5-Cre$^{ERT2}$(PAC)/mTmG mice treated with peanut oil only or with tamoxifen. Cre$^{-}$ mice treated with tamoxifen were tested as controls.

(G) Representative gating strategy for quantification of CD45+ZsGreen+ cells in mouse bone marrow from Cdh5-CreERT2(BAC)/ZsGreen mice treated with peanut oil only or with tamoxifen. Cre- mice treated with tamoxifen were tested as controls.

### Figure S2

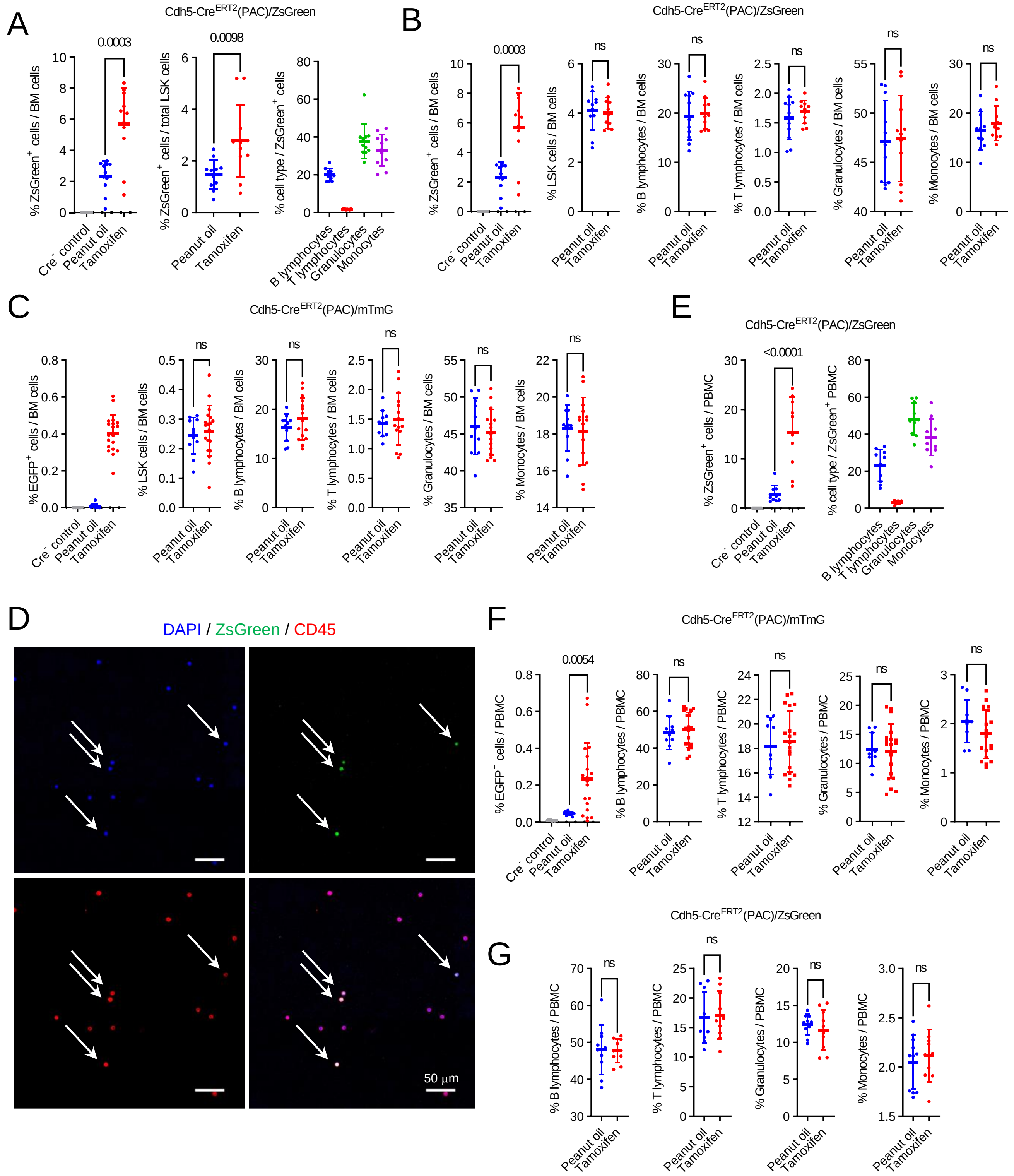

A
Cdh5-CreERT2(PAC)/ZsGreen
0.0003
% ZsGreen+ cells / BM cells
Cre- control
Peanut oil
Tamoxifen
0.0098
% ZsGreen+ cells / total LSK cells
% cell type / ZsGreen+ cells
B lymphocytes
T lymphocytes
Granulocytes
Monocytes
B
Cdh5-CreERT2(PAC)/ZsGreen
0.0003
ns
% LSK cells / BM cells
% B lymphocytes / BM cells
% T lymphocytes / BM cells
% Granulocytes / BM cells
% Monocytes / BM cells
C
Cdh5-CreERT2(PAC)/mTmG
% EGFP+ cells / BM cells
E
Cdh5-CreERT2(PAC)/ZsGreen
<0.0001
% ZsGreen+ cells / PBMC
% cell type / ZsGreen+ PBMC
D
DAPI / ZsGreen / CD45
50 µm
F
Cdh5-CreERT2(PAC)/mTmG
0.0054
% EGFP+ cells / PBMC
% B lymphocytes / PBMC
% T lymphocytes / PBMC
% Granulocytes / PBMC
% Monocytes / PBMC
G
Cdh5-CreERT2(PAC)/ZsGreen

**Figure S2 related to Figure 1. Characterization of tracked hematopoietic progenitors and mature cells in adult bone marrow and peripheral blood of Cdh5-Cre reporter mice.**

(A) Percent ZsGreen$^{+}$ cells in the bone marrow (BM) of Cdh5-Cre$^{ERT2}$(PAC)/ZsGreen mice treated or not treated with tamoxifen showing tamoxifen-independent ZsGreen fluorescence and the significant increase of ZsGreen fluorescence induced by tamoxifen. Cre$^{-}$ mice treated with tamoxifen were used as controls (left). Distribution of viable BM ZsGreen$^{+}$ cells within LSK (middle). Percent B lymphocytes, T lymphocytes, granulocytes, and monocytes in all ZsGreen$^{+}$ cells (right). Each dot reflects results from individual mice (1 femur plus 1 tibia combined); the group means ± SD (error bars) are shown. *P* values (unpaired two-tailed Student's *t*-test) are shown.

(B) Left panel: percent ZsGreen$^{+}$ cells in the bone marrow of Cdh5-Cre$^{ERT2}$(PAC)/ZsGreen mice treated or not treated with tamoxifen. All other panels: the overall (ZsGreen$^{+}$ and ZsGreen$^{-}$) percent LSK cells, B lymphocytes, T lymphocytes, granulocytes and monocytes is similar in the bone marrow of control Cre$^{+}$ mice (peanut oil treated, no tamoxifen) and tamoxifen treated Cre$^{+}$ mice. This indicates that tamoxifen administration has not altered the distribution of bone marrow cell subsets analyzed. Results from individual mice are displayed as dots; means (± SD) are displayed as horizontal lines and error bars. Statistical significance of differences determined by unpaired two-tailed Student's *t*-test are shown; $P<0.05$ is considered statistically significant; ns: not significant difference.

(C) Left panel: percent EGFP$^{+}$ cells in the bone marrow of Cdh5-Cre$^{ERT2}$(PAC)/mTmG mice four weeks after administration of tamoxifen or peanut oil only. Cre$^{-}$ mice treated with tamoxifen were used as controls. All other panels: the overall (EGFP$^{+}$ and EGFP$^{-}$) percent LSK (Lin$^{-}$, Sca1$^{+}$cKit$^{+}$), B (CD19$^{+}$) and T (CD3$^{+}$) lymphocytes, granulocytes (CD19$^{-}$, CD3$^{-}$, Ly6G$^{+}$, CD11b$^{+}$), and monocytes (CD19$^{-}$, CD3$^{-}$, Ly6G$^{-}$, CD11b$^{+}$) is similar in the bone marrow of control Cre$^{+}$ mice (peanut oil treated, no tamoxifen) and tamoxifen treated Cre$^{+}$ mice. Results from individual mice are displayed as dots; means (± SD) are displayed as horizontal lines and error bars. Statistical significance of differences determined by unpaired two-tailed Student's *t*-test are shown; ns: not significant difference.

(D) Representative immunofluorescence image of peripheral blood ZsGreen$^{+}$CD45$^{+}$ nucleated (DAPI$^{+}$) cells (pointed by the arrows) from a tamoxifen treated Cdh5-Cre$^{ERT2}$(PAC)/ZsGreen mouse.

(E) Percent ZsGreen$^{+}$ cells within PBMC of Cdh5-Cre$^{ERT2}$(PAC)/ZsGreen$^{+}$ mice treated with tamoxifen, peanut oil (no tamoxifen) or Cre$^{-}$ mice treated with tamoxifen (left), and relative contribution of B lymphocytes, T lymphocytes, granulocytes, and monocytes to ZsGreen$^{+}$ PBMC (right).

(F) Left panel: percent EGFP$^{+}$ cells within peripheral blood mononuclear cells (PBMC) of Cdh5-Cre$^{ERT2}$(PAC)/mTmG mice treated with tamoxifen or with peanut oil (no tamoxifen). Cre$^{-}$ mice treated with tamoxifen are included as controls (left). All other panels: similar percent B lymphocytes (CD19$^{+}$), T lymphocytes (CD3$^{+}$), granulocytes (CD19$^{-}$, CD3$^{-}$, Ly6G$^{+}$, CD11b$^{+}$), and monocytes (CD19$^{-}$CD3$^{-}$Ly6G$^{-}$CD11b$^{+}$) is detected in PBMC of Cdh5-Cre$^{ERT2}$(PAC)/mTmG mice treated with peanut oil only (no tamoxifen) or with tamoxifen, showing that tamoxifen administration has not altered the distribution of PBMC subsets analyzed.

(G) Similar percent B lymphocytes (CD19$^{+}$), T lymphocytes (CD3$^{+}$), granulocytes (CD19$^{-}$CD3$^{-}$Ly6G$^{+}$CD11b$^{+}$), and monocytes (CD19$^{-}$CD3$^{-}$Ly6G$^{-}$CD11b$^{+}$) is measured in PBMC of Cdh5-Cre$^{ERT2}$(PAC)/ZsGreen mice treated with peanut oil only (no tamoxifen) or with tamoxifen, showing that tamoxifen administration has not altered the distribution of PMBC subsets analyzed.

### Figure S3

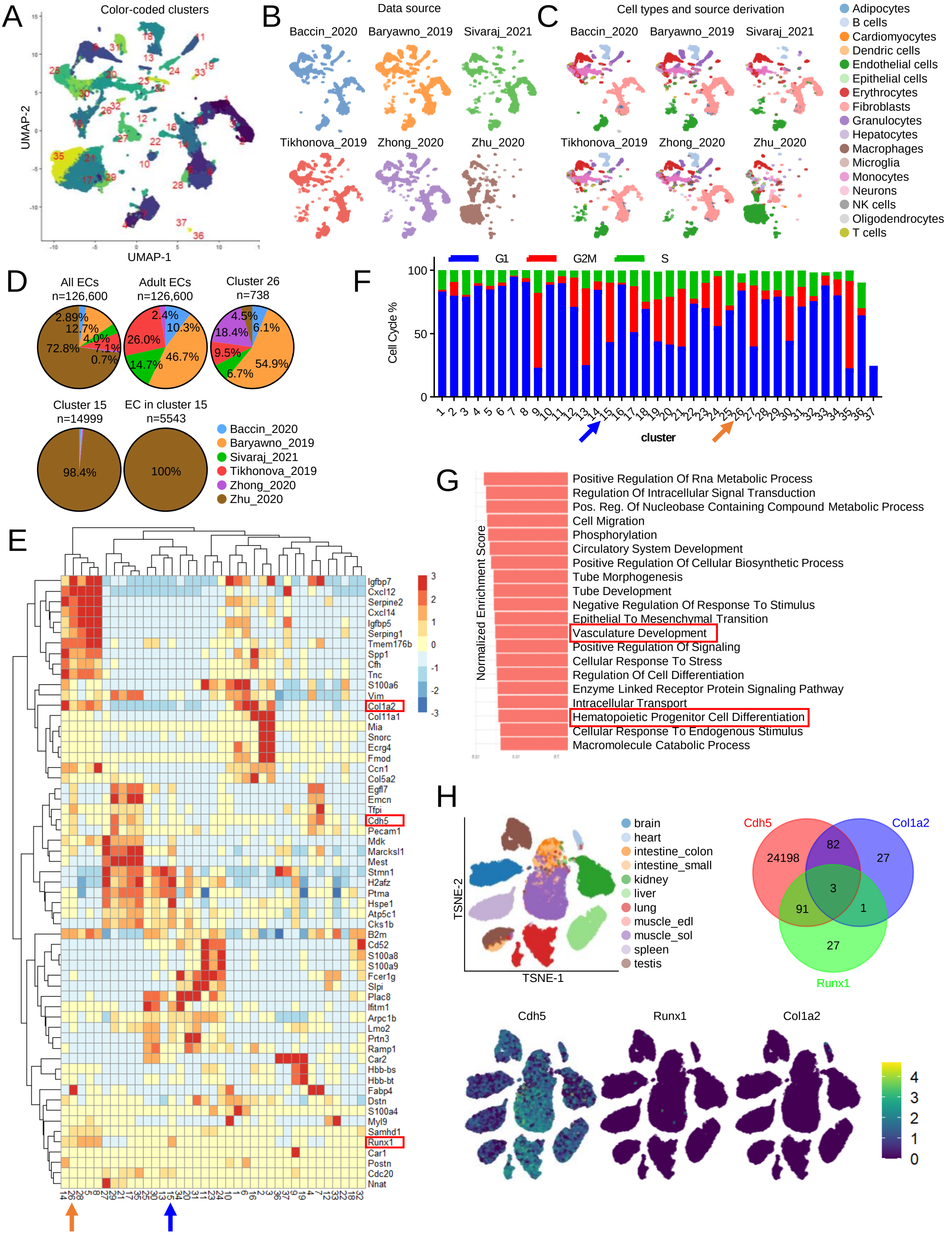
A
Color-coded clusters
UMAP-2
UMAP-1
B
Data source
Baccin_2020
Baryawno_2019
Sivaraj_2021
Tikhonova_2019
Zhong_2020
Zhu_2020
C
Cell types and source derivation
Baccin_2020
Baryawno_2019
Sivaraj_2021
Tikhonova_2019
Zhong_2020
Zhu_2020
Adipocytes
B cells
Cardiomyocytes
Dendric cells
Endothelial cells
Epithelial cells
Erythrocytes
Fibroblasts
Granulocytes
Hepatocytes
Macrophages
Microglia
Monocytes
Neurons
NK cells
Oligodendrocytes
T cells
D
All ECs
n=126,600
2.89%
12.7%
4.0%
7.1%
0.7%
72.8%
Adult ECs
n=126,600
2.4%
10.3%
26.0%
46.7%
14.7%
Cluster 26
n=738
4.5%
6.1%
18.4%
9.5%
54.9%
6.7%
Cluster 15
n=14999
98.4%
EC in cluster 15
n=5543
100%
Baccin_2020
Baryawno_2019
Sivaraj_2021
Tikhonova_2019
Zhong_2020
Zhu_2020
E
Igfbp7
Cxcl12
Serpine2
Cxcl14
Igfbp5
Serping1
Tmem176b
Spp1
Cfh
Tnc
S100a6
Vim
Col1a2
Col11a1
Mia
Snorc
Ecrg4
Fmod
Ccn1
Col5a2
Egfl7
Emcn
Tfpi
Cdh5
Pecam1
Mdk
Marcksl1
Mest
Stmn1
H2afz
Ptma
Hspe1
Atp5c1
Cks1b
B2m
Cd52
S100a8
S100a9
Fcer1g
Slpi
Plac8
Ifitm1
Arpc1b
Lmo2
Prtn3
Ramp1
Car2
Hbb-bs
Hbb-bt
Fabp4
Dstn
S100a4
Myl9
Samhd1
Runx1
Car1
Postn
Cdc20
Nnat
3
2
1
0
-1
-2
-3
F
G1
G2M
S
Cell Cycle %
100
50
0
cluster
G
Normalized Enrichment Score
Positive Regulation Of Rna Metabolic Process
Regulation Of Intracellular Signal Transduction
Pos. Reg. Of Nucleobase Containing Compound Metabolic Process
Cell Migration
Phosphorylation
Circulatory System Development
Positive Regulation Of Cellular Biosynthetic Process
Tube Morphogenesis
Tube Development
Negative Regulation Of Response To Stimulus
Epithelial To Mesenchymal Transition
Vasculature Development
Positive Regulation Of Signaling
Cellular Response To Stress
Regulation Of Cell Differentiation
Enzyme Linked Receptor Protein Signaling Pathway
Intracellular Transport
Hematopoietic Progenitor Cell Differentiation
Cellular Response To Endogenous Stimulus
Macromolecule Catabolic Process
H
TSNE-2
TSNE-1
brain
heart
intestine_colon
intestine_small
kidney
liver
lung
muscle_edl
muscle_sol
spleen
testis
Cdh5
Col1a2
Runx1
24198
82
27
3
91
1
27
Cdh5
Runx1
Col1a2
4
3
2
1
0

#### Figure S3 related to Figure 2. Bioinformatic identification and characterization of prospective Runx1$^+$ hemogenic endothelial cells in adult mouse bone marrow.

(A) Identification of cell clusters 26 and 15 by Uniform Manifold Approximation and Projection (UMAP) visualization; the results reflect color coded unsupervised clustering of six datasets by the Louvain method.

(B) Illustration of the contribution of each of the six datasets to the overall cluster identification. The source of each dataset is identified by color and the relative contribution of each dataset to the overall cell clustering is shown.

(C) Cell type annotation is color-coded in each dataset.

(D) Data source and source contribution to all endothelial cells, adult endothelial cells, and endothelial clusters 15 and 26.

(E) Heatmap depiction of marker genes (top 15) most discriminating each cluster showing that cells in cluster 26 co-express endothelial and mesenchymal markers and the transcription factor Runx1. The orange arrow points to cluster 26; the blue arrow points to cluster 15.

(F) Cell cycle distribution of cells within each cluster showing that cluster 26 cells mostly reside in G1 whereas cells of cluster 15 mostly reside in G2M and S. Cell cycle distribution was assessed by SingleR package as detailed in the methods section.

(G) Gene set enrichment analysis (GSEA) shows that cells in cluster 26 display a selective enrichment in expression of gene sets defining Vascular Development and Hematopoietic Progenitor Cell Differentiation compared to the other clusters as reflected by the normalized enrichment score.

(H) Dataset source: single cell transcriptome atlas of murine endothelial cells from the 11 indicated tissues. Identification of cell clusters by t-distributed stochastic neighbor embedding (t-SNE) visualization. The results are shown as color-coded representation of tissue derivation (top left) and expression levels of Cdh5, Runx1 and Col1a2. The Venn diagram shows that co-expression of Cdh5, Col1a2 and Runx1 is rare in the dataset.

### Figure S4

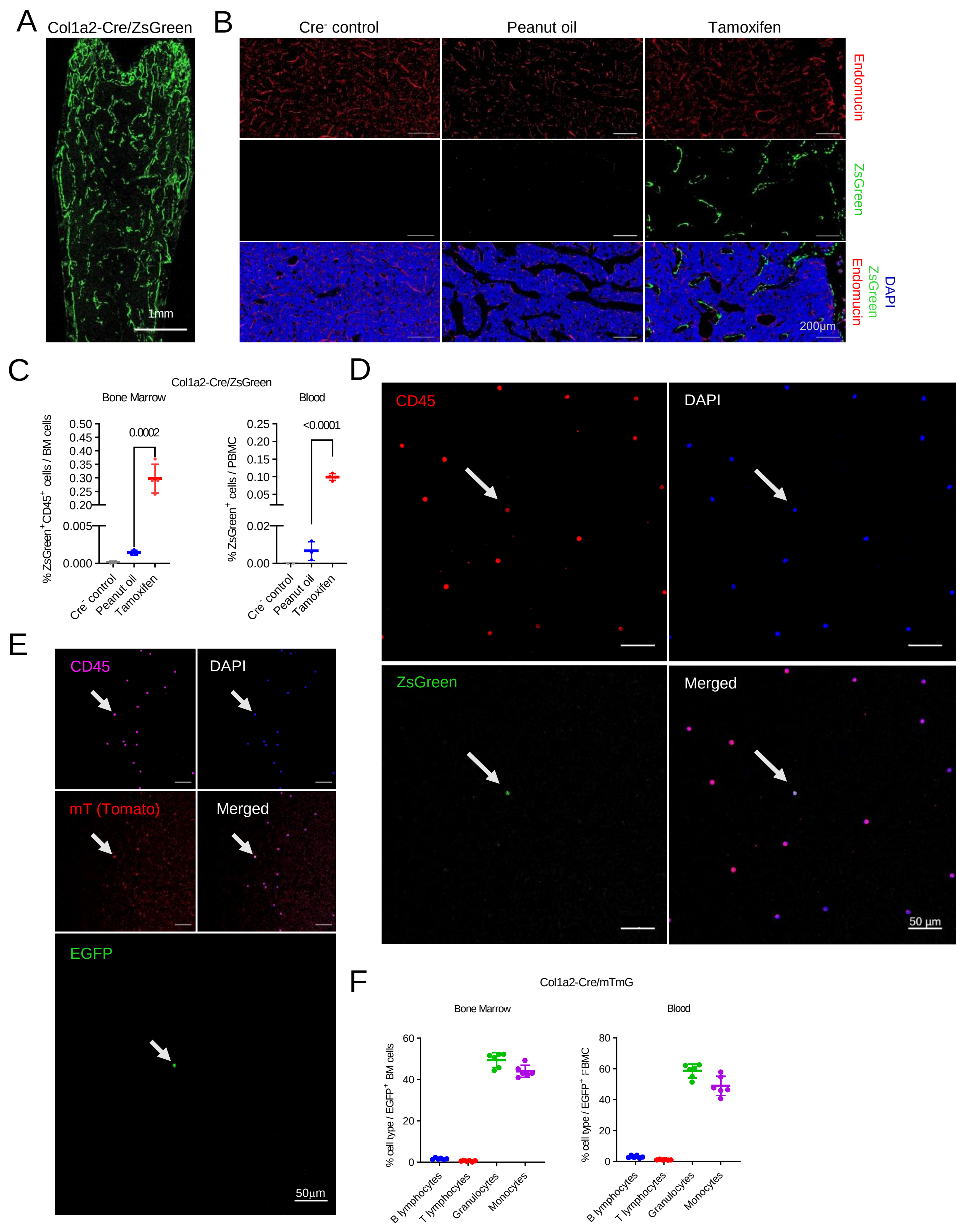
A
Col1a2-Cre/ZsGreen
1mm
B
Cre- control
Peanut oil
Tamoxifen
Endomucin
ZsGreen
DAPI
200µm
C
Col1a2-Cre/ZsGreen
Bone Marrow
Blood
% ZsGreen+CD45+ cells / BM cells
% ZsGreen+ cells / PBMC
0.0002
<0.0001
Cre- control
Peanut oil
Tamoxifen
D
CD45
DAPI
ZsGreen
Merged
50 µm
E
CD45
DAPI
mT (Tomato)
Merged
EGFP
50µm
F
Col1a2-Cre/mTmG
Bone Marrow
Blood
% cell type / EGFP+ BM cells
% cell type / EGFP+ PBMC
B lymphocytes
T lymphocytes
Granulocytes
Monocytes

**Figure S4 related to Figure 3. Characterization of Col1a2-tracked cell populations in bone marrow and blood.**

(A) Representative confocal image of bone marrow from a Col1a2-Cre/ZsGreen mouse induced with tamoxifen showing that the $ZsGreen^{+}$ cells are widely detected.

(B) In the bone marrow of Col1a2-Cre/ZsGreen mice, only few $ZsGreen^{+}$ cells are detected in the bone marrow of a peanut oil-treated mouse (middle panels), whereas numerous $ZsGreen^{+}$ cells are detected in the tamoxifen treated mouse (right panel). No $ZsGreen^{+}$ cells are detected in a $Cre^{-}$ control bone marrow (left panels). Representative images.

(C) Percent $ZsGreen^{+}$ cells among bone marrow $CD45^{+}$ cells (left) and peripheral blood mononuclear cells (PBMC, right) detected in Col1a2-$Cre^{ERT2}$/ZsGreen mice after peanut oil or tamoxifen treatment. Each dot represents results from individual mice (1 femur plus 1 tibia combined). The means (± SD) are displayed as horizontal lines and error bars. Statistical significance of differences determined by unpaired two-tailed Student's *t*-test are shown.

(D) Representative confocal image of a blood smear from a Col1a2-Cre/ZsGreen mouse treated with tamoxifen showing the presence of nucleated $CD45^{+}$ cells tracked by ZsGreen/Col1a2 fluorescence (pointed by the arrows).

(E) Representative confocal image of a blood smear from a Col1a2-Cre/mTmG mouse treated with tamoxifen showing the presence of nucleated $CD45^{+}$ cells tracked by EGFP/Col1a2 fluorescence (pointed by the arrows).

(F) Distribution of B and T lymphocytes, granulocytes, and monocytes in EGFP+ bone marrow (BM) and EGFP+ peripheral blood mononuclear cells (PBMC) from Col1a2-Cre/mTmG mice treated with tamoxifen. Each dot represents results from individual mice (1 femur plus 1 tibia combined).
