## Supplemental Table1 for "Bone marrow hemogenic endothelial cells contribute multilineage hematopoietic progenitors in adult mice"

Methods to Culture and/or Immortalize AHEC

The methods below, alone and variously combined, were used unsuccessfully to culture or immortalize sorted AHEC.

| Culture Condition | Culture Surfaces | Lentiviral Constructs |
| --- | --- | --- |
| M199 (with Earle’s salts and L-glutamine), L-Ascorbic Acid, Sodium Salt Heparin, Sodium Salt Glutamine, EC growth factor bovine – Sigma Cat#E-9640 (100x), heat inactivated mouse serum.(Hashida et al., 2002) | gelatin coated plates or flasks | myr-HA-Akt1 |
| Non-essential amino acid, 10mM Hepes, 100ug/mL Heparin, 50ug/mL endothelial cell growth supplement,  20% FBS in DMEM low + Hams F12 Media.(Poulos et al., 2015) | FN coated plates or flasks | polyoma middle T-antigen |
| Co-cultured with OP9 stromal cells. RPMI, supplemented with 10% FCS and 10-5 mol/L 2-mercaptoethanol. VEGF (10 ng/mL) was added every three days.(Wakabayashi et al., 2018) | Corning® ECM surfaces | mCdk4 |
| IMDM medium containing 15% FCS, 1% L-glutamine, 1% BSA, 10^-4^ mM 2-mercaptoethanol, 0.01 mg/ml rh insulin, 100 ng/ml rm VEGF, 100 ng/ml rm bFGF, and 100 ng/ml rm, and 0.2 mg/ml human transferring.(Fang et al., 2012) | Corning® Primaria™ Surface | mTert |
| IMDM medium containing 15% FBS, 1% BSA, 10^−4^ mM 2-mercaptoethanol, 1% Glutamax, 1% ITS-X, 100 ng/ml rmVEGFa and 50 ng/ml rmbFGF. (Yu et al., 2016) | Poly-L-Lysine cultureware (Corning) | mCdk4 + mTert |
| Semi-solid methylcellulose (Stem Cell Technologies), supplemented with 100 ng/ml mVEGFa.(Yu *et al.*, 2016) | Poly-D-Lysine cultureware (Corning) |  |
| EGM™-2 Endothelial Cell Growth Medium-2 BulletKit(Lonza, # CC-3162) | Collagen I / Collagen II coated plates or flasks |  |
| EGMTM -2 MV Microvascular Endothelial Cell Growth Medium-2 BulletKitTM (Lonza, # CC-3202) | Laminin coated plates or flasks |  |
| MethoCult (StemCell, #GF3434) |  |  |
